# Partitioning plant spectral diversity into alpha and beta components

**DOI:** 10.1101/742080

**Authors:** Etienne Laliberté, Anna K. Schweiger, Pierre Legendre

## Abstract

Plant spectral diversity — how plants differentially interact with solar radiation — is an integrator of plant chemical, structural, and taxonomic diversity that can be remotely sensed. We propose to measure spectral diversity as spectral variance, which allows the partitioning of the spectral diversity of a region, called spectral *gamma* (*γ*) diversity, into additive alpha (*α*; within communities) and beta (*β*; among communities) components. Our method calculates the contributions of individual bands or spectral features to spectral *γ*-, *β*-, and *α*-diversity, as well as the contributions of individual plant communities to spectral diversity. We present two case studies illustrating how our approach can identify “hotspots” of spectral *α*-diversity within a region, and discover spectrally unique areas that contribute strongly to *β*-diversity. Partitioning spectral diversity and mapping its spatial components has many applications for conservation since high local diversity and distinctiveness in composition are two key criteria used to determine the ecological value of ecosystems.

## INTRODUCTION

Major environmental changes, including land-use change, climate change, and invasive species are altering the Earth’s biodiversity. The rapid rate and broad extent of those changes far exceed our capacity to monitor them via field-based sampling alone. This calls for the development of new remote sensing approaches that can provide rapid estimates of biodiversity over broad regions (Pereira *et al.* 2013; Turner 2014; Bush *et al.* 2017). For terrestrial plants, imaging spectroscopy is emerging as the most promising remote sensing method for estimating biodiversity (Féret & Asner 2014; Wang & Gamon 2019). This is because its high spectral resolution allows plant species to be discriminated from one another, while also enabling the determination of ecologically important foliar functional traits (Asner & Martin 2009; Ustin *et al.* 2009).

For every pixel of an aerial image, imaging spectroscopy measures reflected solar radiation in tens to hundreds of contiguous, narrow (~10 nm wide) wavelength bands, usually covering all or part of the visible to shortwave infrared range (400–2500 nm) of the electromagnetic spectrum. Leaf “spectral signatures” of plants provide unique expressions among species of how solar radiation interacts with photosynthetic pigments, water, proteins, as well as structural and chemical defense compounds, and thus represent the evolution of plant adaptations to different environmental conditions (Cavender-Bares *et al.* 2016; McManus *et al.* 2016). At the crown scale, these spectral signatures are further influenced by architectural traits due to scattering of photons within canopies (Asner 1998; Ollinger 2010). Therefore, plant spectral diversity is emerging as an integrator of plant chemical, structural and taxonomic diversity that can be remotely sensed (Cavender-Bares *et al.* 2017; Schweiger *et al.* 2018; see also Appendix S1 in Supporting Information).

One of the most influential conceptual developments in community ecology has been Whittaker’s (1960, 1972) suggestion to partition biodiversity across space into *α*, *β*, and *γ* components. Originally, *α* diversity was defined as the species diversity *within* communities, and *β* as the variation in species composition *among* communities; together, *α*- and *β*-diversities jointly determined *γ*-diversity, which is the species diversity across an entire region of interest. In this paper, we transpose this foundational ecological concept from species diversity to spectral diversity (Fig. 1). This requires that the spatial resolution of the imagery matches the size of the object of interest (Woodcock & Strahler 1987), meaning that pixels should be approximately equal or smaller than the size of an average canopy plant. At such fine spatial resolutions, the relationship between spectral and taxonomic diversity is strongest (Wang *et al.* 2018a) and imaging spectroscopy can provide direct, spatially explicit estimates of plant alpha (*α*; within community) diversity (Féret & Asner 2014; Wang *et al.* 2018b), and can detect changes in plant community composition across landscapes (Draper *et al.* 2019). The ability to generate wall-to-wall, high-resolution maps of canopy plant diversity across entire regions brings tremendous benefits for biodiversity science and conservation (e.g., Asner *et al.* 2017); however, conceptual and methodological challenges remain, especially with regard to *β*-diversity estimation (Rocchini *et al.* 2010, 2018).

**Figure 1.**
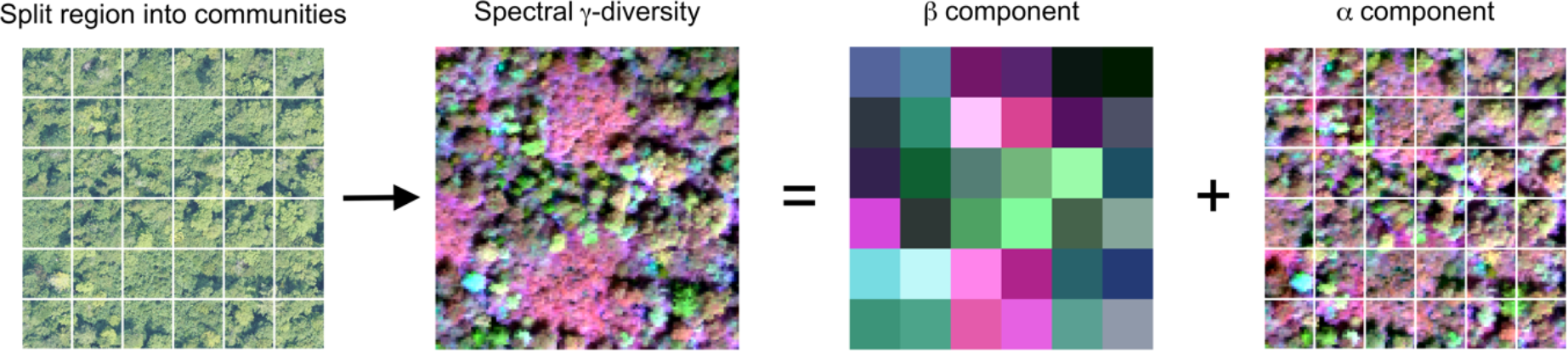
Partitioning plant spectral γ-diversity into additive *β* and *α* components. A region of interest is split into a number of communities of a specific size and shape (here, 20 m × 20 m squares, representing standard forest inventory plots). Spectral *γ*-diversity refers to the total spectral diversity in the entire region calculated from pixel-level reflectance. The *β* component corresponds to spectral diversity *among* communities, with similar colours sharing more similar spectral composition. The *α* component refers to spectral diversity *within* individual communities. The left-most panel is a true colour (red-green-blue, RGB) image of an area of Bartlett Experimental Forest; colours for the other panels were obtained using the reflectance of different wavelength bands (R = 779 nm, G = 639 nm, B = 2301 nm), followed by linear stretching.

Spectral diversity is sometimes called spectral heterogeneity or spectral variability (Rocchini *et al.* 2010), and has been defined as spatial variation in spectral reflectance (Rocchini *et al.* 2010; Ustin & Gamon 2010; Gholizadeh *et al.* 2018; Wang & Gamon 2019). Intuitively, spectral diversity can be conceptualized as multivariate dispersion, for which there are various statistical measures highlighting different aspects of spectral diversity. For example, Wang et al. (2018a) used the average coefficient of variation (CV) of each band for a set of pixels, whereas Rocchini et al. (2010) used the mean distance from the spectral centroid; we note that the latter has also been proposed as a measure of functional diversity in multivariate trait space (Laliberté & Legendre 2010). However, none of the currently used metrics allow the partitioning of spectral diversity into its *α* (within communities) and *β* (among communities) components (Fig. 1).

Here we propose to use the *spectral variance* among image pixels as a measure of spectral diversity. Our approach builds on that of Legendre and De Cáceres (2013) for species inventory data, adapts it to spectral data, and extends it to jointly consider *α*-, *β*-, and *γ*-diversity. Casting spectral diversity as spectral variance has a number of benefits:

1. the classical partitioning of sums of squares allows us to partition spectral *γ*-diversity into additive spectral *α*- and *β*-diversity components (Fig. 1), from which the relative importance of local and regional processes regulating spectral diversity across a region of interest can be inferred;
2. it allows us to estimate the contributions of individual plots or communities to spectral *β*-diversity, highlighting areas that are spectrally distinct within the broader region;
3. it allows us to calculate the contributions of individual bands or spectral features to spectral *γ*-, *β*- or *α*-diversity (Fig. 2), providing information about the underlying biological traits driving spectral diversity;
4. it is easily implemented in software packages in a computationally efficient way, which is important when dealing with high-volume image data;
5. it provides a direct link to other statistical procedures based on least squares (e.g., MANOVA, multiple linear regression, canonical redundancy analysis, *K*-means partitioning).

**Figure 2.**
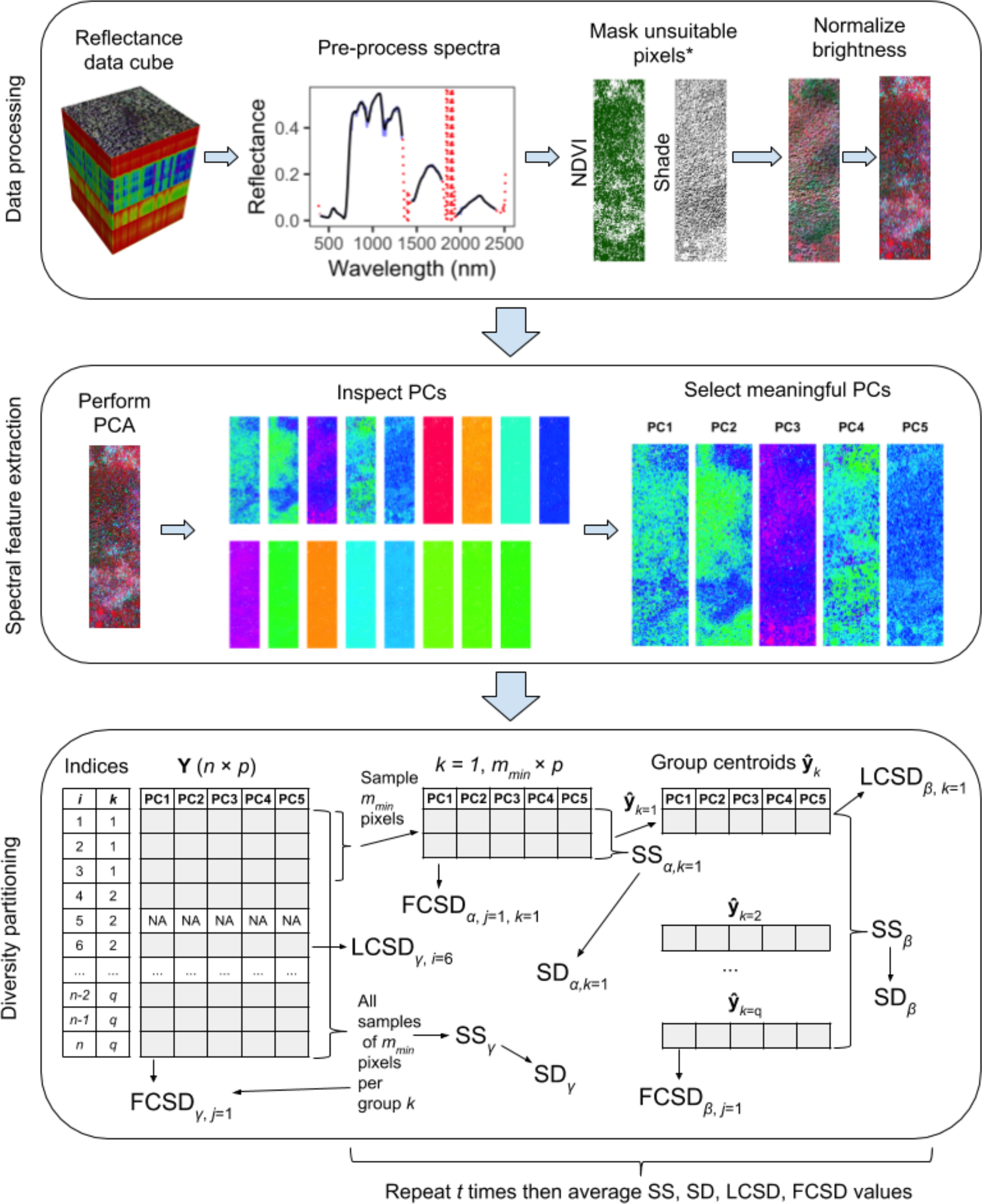
Overview of our proposed workflow for partitioning plant spectral diversity. In our NEON case study, spectral data pre-processing included removing atmospheric water absorption bands (wavelengths between 1340–1455 nm and 1790–1955 nm) and noisy regions of the spectrum (wavelengths <400 nm and >2400 nm), and applying a Savitzky-Golay filter (order = 3, size = 7) to every pixel in the image to remove high-frequency noise. We masked all pixels with normalized difference vegetation index (NDVI) values <0.8, and brightness-normalized all spectra (Feilhauer *et al.* 2010). Then, we performed a PCA with type I-scaling, and visually inspected the first 17 PCs which together accounted for >99% of the total spectral variation among all pixels (Fig. S3). Only the first five PCs showed meaningful biological spatial patterns and were retained for spectral diversity measurements; PCs 6–17 were excluded based on visual inspection as they expressed artefacts from image acquisition and processing (Fig. S3). For illustration purposes, in the diversity partitioning analysis (bottom panel) we show communities composed of only three pixels, whereas in fact we used a community size of 40 × 40 pixels in our NEON case study. For abbreviations see text. *The shade mask is for illustrative purposes and is not applied to the PCs shown in the middle panel.

After describing the theory behind our spectral diversity partitioning approach, we illustrate it using a simulation. We then apply our method to imaging spectroscopy data collected over the Bartlett Experimental Forest by the National Ecological Observatory Network (NEON) Airborne Observation Platform (AOP; Kampe *et al.* 2010). The R code and data for our analyses are available online (https://github.com/elaliberte/specdiv).

## PARTITIONING SPECTRAL DIVERSITY

### Size and shape of spatial units

Partitioning spectral *γ*-diversity into its *α* and *β* components first requires defining the extent of the region of interest (Fig. 1). Delineating the region of interest is relatively straightforward since it corresponds to the region over which imagery is acquired or a subset thereof. Delineating the size and shape of communities across the region of interest, however, is more difficult. What constitutes an ecological community has been the subject of considerable debate (see review by Ricklefs 2008). Generally, a community is defined as “a group of organisms representing multiple species living in a specified place and time” (Vellend 2010). This definition implies that a community must be larger than the size of an individual organism, but how much larger will depend on the objectives of the study. For the purpose of this work, we focus on communities of canopy plants, because these are the organisms that can be seen in aerial images. We use “community” in the sense of “sampling unit” in vegetation surveys, which can be defined as the area in which the species composition of the vegetation type of interest is adequately represented (Mueller-Dombois & Ellenberg 1974).

Setting the size of a community to the size of typical inventory plot for a given ecosystem type facilitates interpretation as this is the sampling unit that field ecologists are familiar with. For example, forest inventory plots often measure 20 m × 20 m (Fig. 1), which is large enough to include several canopy trees. However, we recognize that setting fixed and regularly shaped boundaries to delineate communities is artificial (Ricklefs 2008), and point out that community size and shape can be changed in the analysis.

### Spectral gamma (*γ*) diversity

Let **Y** = [*y*_*ij*_] be a matrix containing the positions, along the *p* axes defining the *spectral space* (column vectors **y**_1_, **y**_2_, … **y**_*p*_ of **Y**), of *n* pixels (row vectors **x**_1_, **x**_2_, … **x**_*n*_ of **Y**) in a region of interest (Fig. 2). We use indices *i* and *j* to denote rows (pixels) and columns (axes) of matrix **Y**, respectively. The *p* axes could be all or a subset of the original spectral bands, a set of vegetation indices calculated from selected spectral bands (Bannari *et al.* 1995), or a set of *p* uncorrelated spectral features extracted using dimensionality reduction methods such as principal component analysis (PCA). We use PCA in this section and in our case studies and point out the mathematical relationships between the principal components (PCs) and spectral variation below. We use the general term *variation* for sums of squares (SS, an abbreviation for “sum of the squared deviations from the mean”), and reserve the term *variance* when talking about spectral diversity (SD).

We refer to the total spectral diversity of the entire region as spectral *γ*-diversity (SD_*γ*_). SD_*γ*_ is measured by the total variance of **Y**, or Var(**Y**). This is done by first computing for every pixel and spectral feature *y*_*ij*_ the squared deviations s_*ij*_ from the average pixel (across the whole region) in terms of spectral reflectance, i.e. the column means of **Y**:

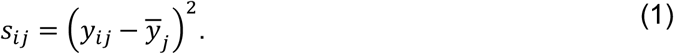

The total sum of squares (SS) of matrix **Y** is calculated by summing all *s*_*ij*_:

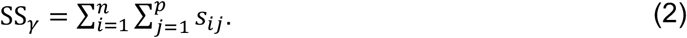

Contrary to SS_*γ*_, SD_*γ*_ is scaled by the number of pixels in the region, such that SD_*γ*_ of regions containing different numbers of pixels can be compared with one another:

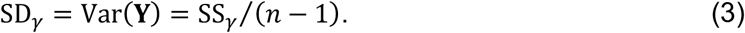

We note that for calculating the joint SD_*γ*_ of adjacent regions, their SS_*γ*_ statistics can be added and divided by the total number of pixels minus one, but their region-level SD_*γ*_ statistics cannot be added directly.

One might be interested in determining the individual contribution of the *j*th spectral feature to SS_*γ*_. We call this the *feature contribution to spectral γ-diversit*y or FCSD_*γ,j*_ (Fig. 2), which can be calculated from the sum of squares of the *j*th feature:

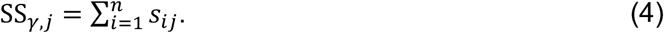

Dividing SS_*γ,j*_ by (*n* − 1) gives the variance of the *j*th feature, or Var(**y**_*j*_). FCSD_*γ,j*_ can then be calculated as:

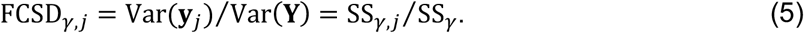

If the *p* features are principal components from PCA scaling type 1, then the FCSD_*γ,j*_ values correspond to their relative eigenvalues. We note that FCSD_*γ,j*_ cannot be mapped because the contribution of each spectral feature applies to the region as a whole.

Likewise, one might wish to estimate the individual contribution of the *i*th pixel within the region to SD_*γ*_. We refer to this as the *local contribution to spectral γ-diversit*y, or LCSD_*γ,i*_, which is calculated as:

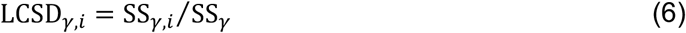

where

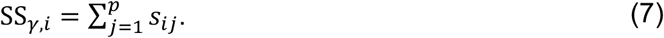

We note that LCSD_*γ,i*_ indices are important visual elements in PCA ordination: each LCSD_*γ,i*_ value corresponds to the squared distance from one pixel to the centroid in the *p*-dimensional PCA ordination plot. In addition, the LCSD_*γ,i*_ can be plotted on maps since one value is associated with every pixel in the image. Doing so indicates which pixels are most spectrally dissimilar from the mean pixel of the region in spectral feature space. We note that the SS_*γ*,*i*_ and LCSD_*γ*,*i*_ indices are additive. The indices from adjacent pixels within an area of interest, for example an individual tree, can be added up, such that their sums represent the local contributions of the area of interest to SS_*γ*_ and SD_*γ*_. LCSD indices are also useful when computed at the community scale (i.e. LCSD_*β*_), because they then correspond to the ecological concept of *β*-diversity; see “Spectral beta (*β*) diversity” below.

### Partitioning the total sum of squares

Partitioning the sum of squares forms the basis of a series of classic statistical approaches based on least squares, such as the analysis of variance (ANOVA). From these methods, it is well known that the total sum of squares of a matrix **Y** (SS_total_) can be partitioned into additive among-group (SS_among_) and within-group (SS_within_) components:

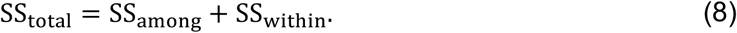

In linear regression analysis, we talk about the SS explained by the regression equation and the residual variation. These two components sum to the total sum of squares.

Using the same indices as in the previous section, the ANOVA relationship can be expressed as:

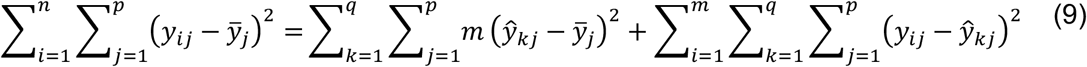

where *q* is the number of groups, *ŷ*_*kj*_ is the arithmetic mean of the *j*th variable (column) for the *k*th group:

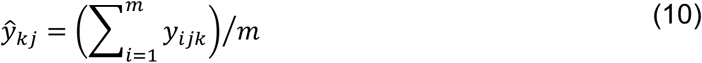

and where *m* is the number of samples (rows) in each group *k*; an important assumption here is that *m* is equal in each group. The proof of this theorem can be found in standard statistics textbooks and is therefore not shown here.

In the next two sections, we apply Equation 9 to partition the total sum of squares of a region SS_*γ*_ into additive among- (*β*) and within-group (*α*) components from which spectral *β*- and *α*-diversity can be calculated directly.

### Spectral beta (*β*) diversity

Let us divide **Y** into *q* groups of *m* spatially contiguous pixels, where each group corresponds to a local community (e.g., a vegetation survey plot); *n* = *q m*. Here, we assume that each of these communities corresponds to a square of equal area, which is 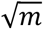 pixels wide; with this setup, each community is represented by the same number of pixels. We will present later our suggestion to use a rarefaction procedure to handle situations where *m* differs among groups.

*Spectral β-diversity*, or SD_*β*_, represents the degree to which the *q* communities within a region differ from each other in terms of spectral composition. We note that SD_*β*_ is a *non-directional* measure of *β-diversity sensu* Anderson et al. (2011). To calculate SD_*β*_, we first compute the squared deviations *s*_*kj*_ of the *k*th community from the average pixel of the region in terms of spectral reflectance, i.e. the column means of **Y** across all variables *j*:

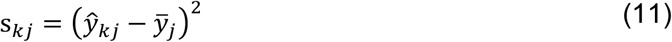

where *ŷ*_*kj*_ is the arithmetic mean of the *k*th community (i.e. the community centroid) for the *j*th spectral feature (Equation 10).

The sum of squares associated with each community *k* (SS_*β,k*_) is:

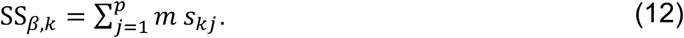

The total sum of squares of the *β* component (SS_*β*_) is:

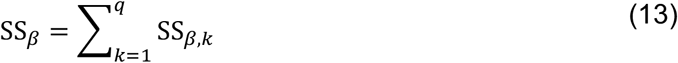

from which SD_*β*_ is calculated as:

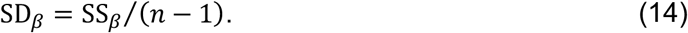

The contribution of each community *k* to SD_*β*_, which we call the *local contribution to spectral β-diversit*y (LCSD_*β,k*_), can be computed by the following ratio of sum of squares:

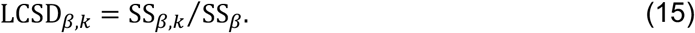

Finally, one can compute the *feature contribution to spectral β-diversit*y or FCSD_*β,j*_ of the *j*th spectral feature as:

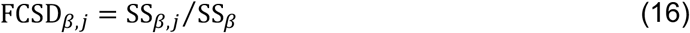

where

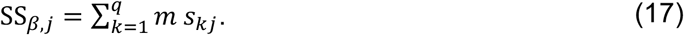

We note here that LCSD_*β,k*_ can be mapped because each community *k* has its own LCSD_*β*_ value. On the other hand, FCSD_*β,j*_ or SD_*β*_ cannot be mapped because they refer to the region as a whole.

### Spectral alpha (*α*) diversity

Spectral *α*-diversity, or SD_*α*_, is the degree to which neighbouring pixels *within* a local community differ spectrally from each other. Contrary to SD_*β*_ and SD_*γ*_, which apply to the entire region, SD_*α*_ is defined at the community level. Therefore, we denote SD_*α*_ by the index *k*, SD_*α,k*_, since it is measured for each community *k*. To calculate SD_*α,k*_, we first compute for every pixel and spectral feature per community *y*_*ijk*_ the squared deviations *s*_*ijk*_ from the mean pixel spectrum of the *k*th community for each spectral feature or column of **Y**:

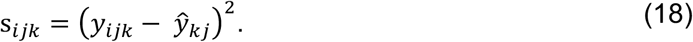

The sum of squares associated with the *j*th spectral feature of community *k* is:

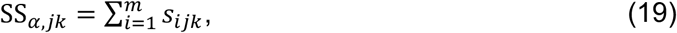

and the total sum of squares for community *k* is:

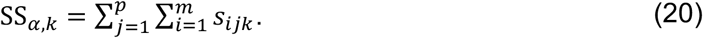

SD_*α,k*_ is obtained by dividing SS_*α,k*_ by (*m* − 1), where *m* is the number of pixels within one community, to make it comparable with other communities differing in their numbers of pixels:

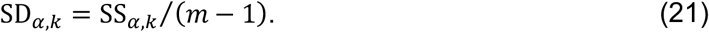

The total sum of squares of the *α*-component for all *q* communities within the entire region is:

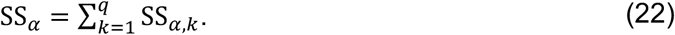

Importantly, following Equations 8 and 9, SS_*α*_ and SS_*β*_ are linked to SS_*γ*_ by the relationship:

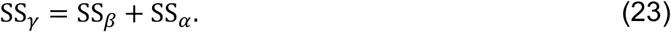

Therefore, SS_*α*_ and SS_*β*_ can be used directly to determine the relative importance of the *α* and *β* components to spectral *γ*-diversity.

Finally, the *contribution of the j*th feature *to the spectral α-diversit*y of the *k*th community, which we call FCSD_*α,jk*_, can be computed as:

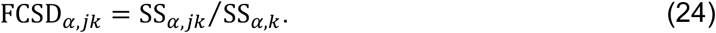

These FCSD_*α,jk*_ values can be mapped and give us useful information about the origin of spectral *α*-diversity across different communities.

## CASE STUDY 1: SIMULATED REGIONS

To illustrate our approach, we first use leaf spectra data to simulate imagery (Appendix S2). This removes much of the complexity associated with real imagery, where one has to deal with much higher numbers of pixels, varying illumination and sensor viewing geometry, and presence of shaded and non-vegetated pixels. We simulated two regions with equal spectral *γ*-diversity, but contrasting spectral *β*- and *α*-diversities (Fig. 3). Each region is composed of 25 × 25 pixels, populated with leaf-level spectra of three temperate tree species (i.e. *Populus deltoides* W. Bartram ex Marshall subsp*. deltoides* Marsh, *P. tremuloides* Michaux, and *Betula alleghaniensis* Britton) measured in the field on 15 individual plants (Fig. S1). These 25 × 25 pixels regions are equally split into 25 communities, each composed of 5 × 5 pixels.

**Figure 3.**
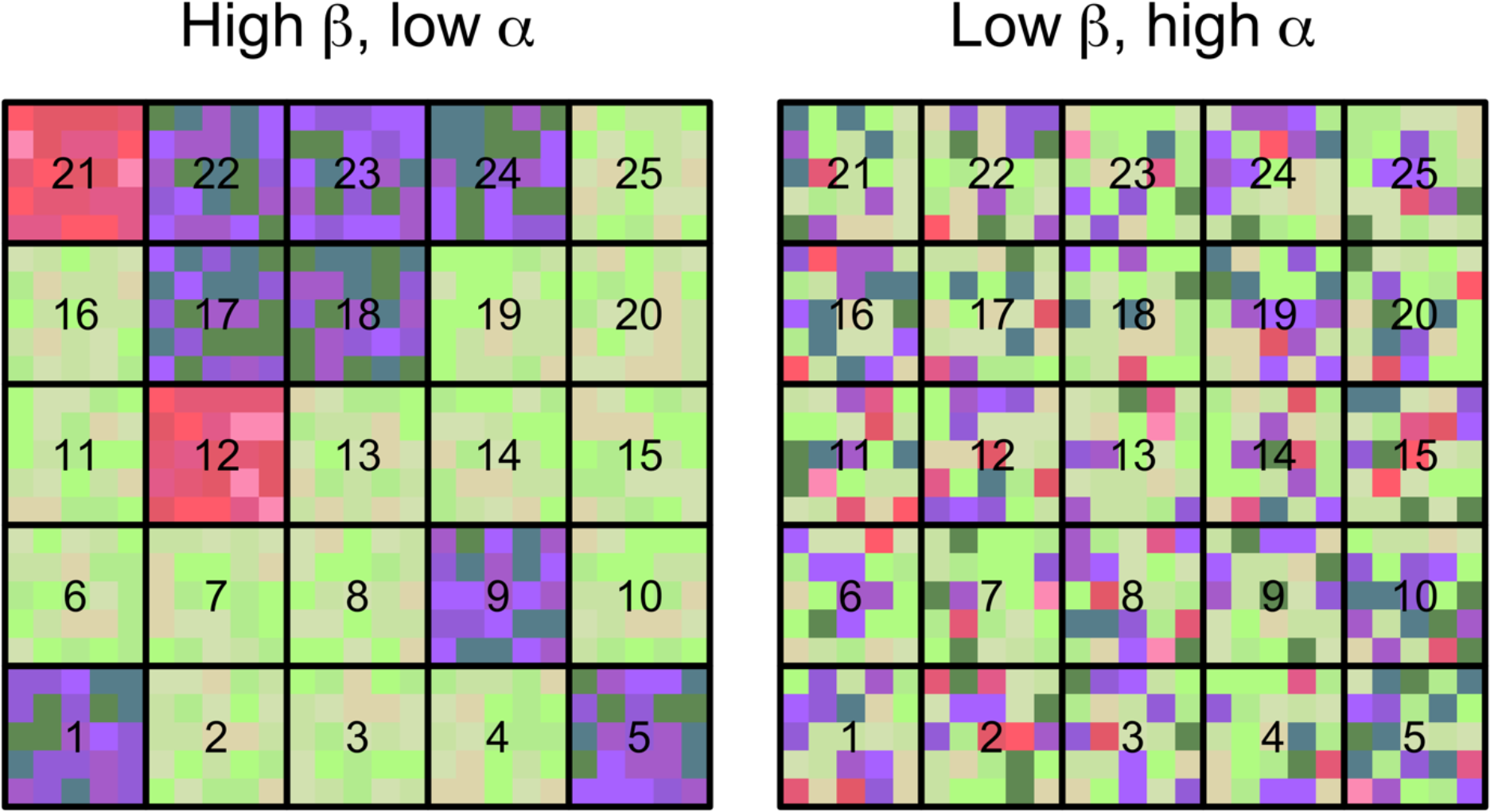
Two simulated landscapes of equal spectral *γ*-diversity, but with contrasting spectral *β*-diversity and *α*-diversity. **Left**: high spectral *β*-diversity but low *α*-diversity. **Right**: low spectral *β*-diversity but high *α*-diversity. Each landscape is composed of 25 communities (numbered black squares), each composed of 5 × 5 pixels (smaller coloured squares). The size of each pixel is equivalent to the size of an individual plant and their colour corresponds to one of the 15 leaf spectra (= 3 species × 5 individuals) shown in Figure S1. These colours were set by mapping the scores of the first three principal components (PC) for each spectrum to a red-green-blue (RGB) scale (PC 1 = green, PC 2 = red, PC 3 = blue). We generated the high spectral *β*-diversity but low spectral *α*-diversity scenario (left panel) by randomly assigning (with replacement) pixels within each community with individual spectra from single species (Fig. S1, bottom row). We selected species identity per community at random using the following probabilities: 0.60 (*Betula alleghaniensis*, green hues), 0.35 (*Populus deltoides*, blue hues) and 0.05 (*Populus tremuloides*, red hues). In this scenario, spectral *β*-diversity was high and spectral *α*-diversity low because interspecific spectral variation (particularly between *Betula* and the two *Populus* species) was higher than intraspecific spectral variation (Fig. S2). Next, to reduce spectral *β*-diversity and increase *α*-diversity while holding *γ*-diversity constant, we moved the pixels of the left panel to randomly selected positions in the right panel.

For both scenarios, we calculated the SS across the entire region (SS_*γ*_), partitioned SS_*γ*_ into its *β* and *α* components, and calculated spectral *γ*- *β-,* and *α*-diversity (Fig. 4a). As spectral features (columns of **Y**) we used the first three PCs (using type-I scaling in PCA), which together explained >97% of the total variation in spectral reflectance. As expected, spectral *γ*-diversity was equal for both scenarios (Table 1), whether expressed as the total sum of squares (SS_*γ*_ = 1.66), or standardized by *n –* 1 pixels (SD_*γ*_ = 0.0027). In addition, in the high *β*-diversity scenario, spectral variation among communities (SS_*β*_, ~84%) largely exceeded spectral variation within communities (SS_*α*_, ~16%), whereas in the low *β*-diversity scenario SS_*β*_ was much lower (~5%) than SS_*α*_ (~95%) (Table 1).

**Table 1.**
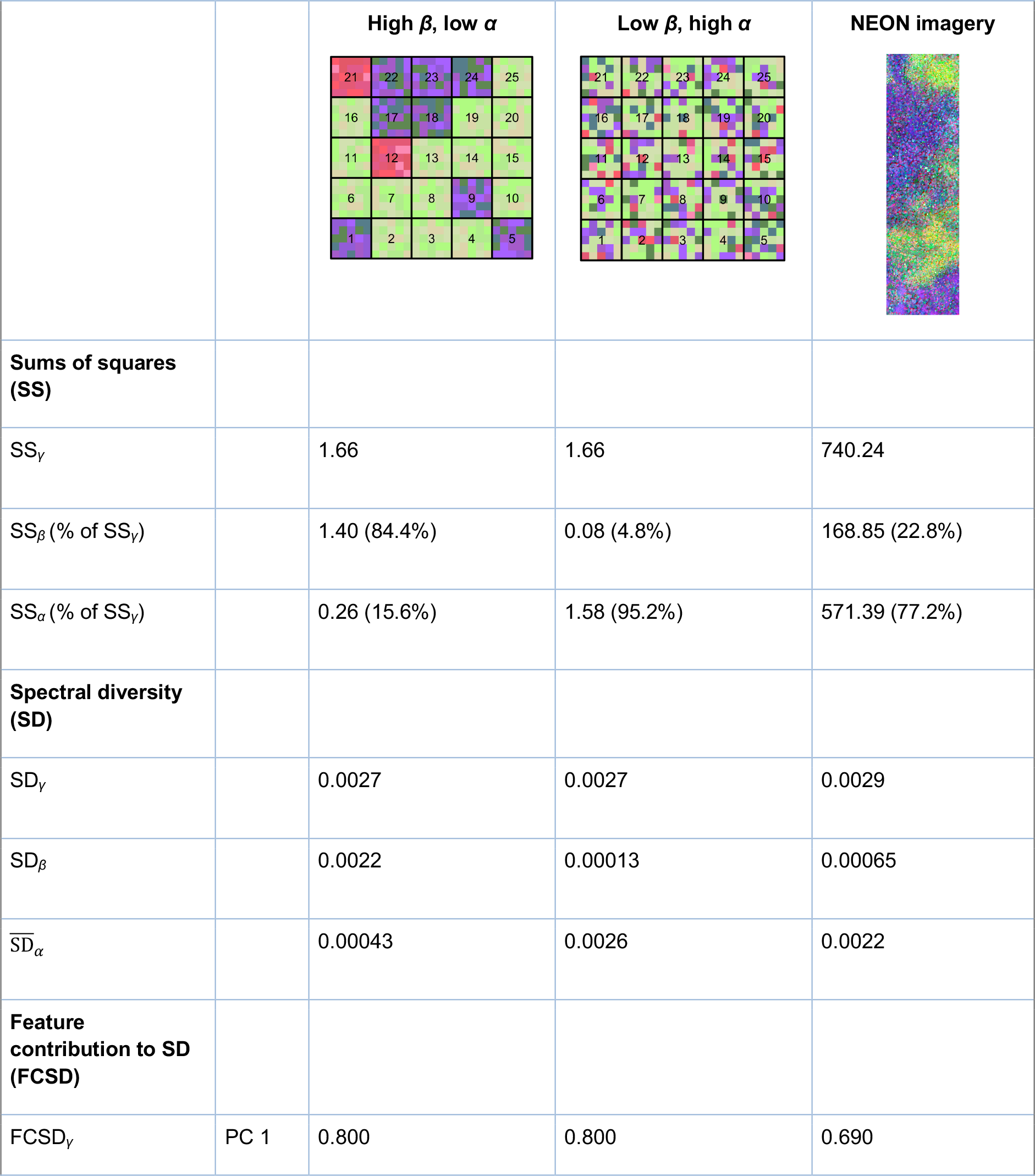
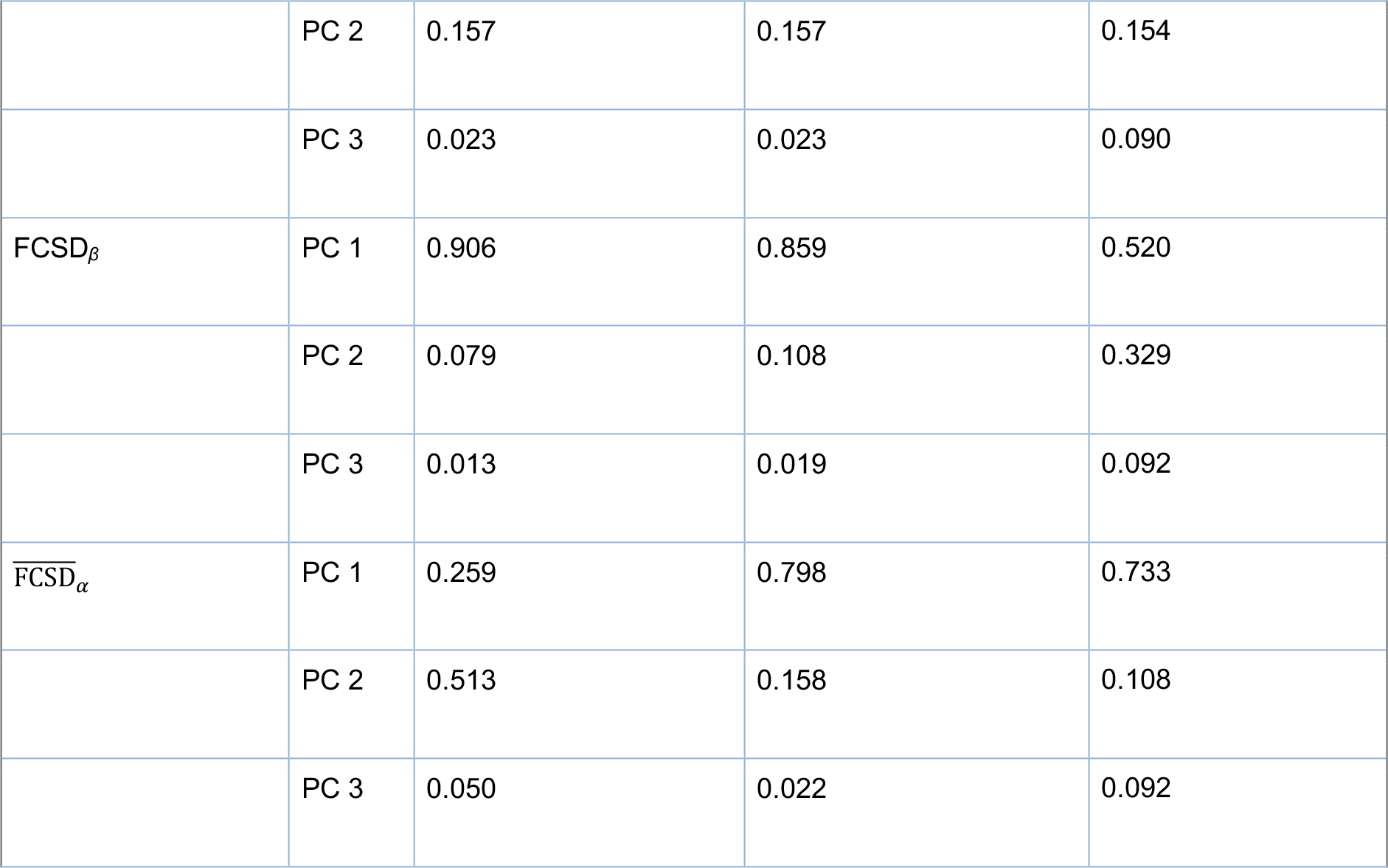
Partitioning spectral diversity into additive components for the two simulated regions (Figs. 3–4) and for the NEON imagery (Fig. 5).

| | | High $\beta$, low $\alpha$ | Low $\beta$, high $\alpha$ | NEON imagery |
| --- | --- | --- | --- | --- |
| **Sums of squares (SS)** |  |  |  |  |
| $SS_Y$ | | 1.66 | 1.66 | 740.24 |
| $SS_\beta$ (% of $SS_Y$) | | 1.40 (84.4%) | 0.08 (4.8%) | 168.85 (22.8%) |
| $SS_\alpha$ (% of $SS_Y$) | | 0.26 (15.6%) | 1.58 (95.2%) | 571.39 (77.2%) |
| **Spectral diversity (SD)** |  |  |  |  |
| $SD_Y$ | | 0.0027 | 0.0027 | 0.0029 |
| $SD_\beta$ | | 0.0022 | 0.00013 | 0.00065 |
| $\overline{SD}_\alpha$ | | 0.00043 | 0.0026 | 0.0022 |
| **Feature contribution to SD (FCSD)** |  |  |  |  |
| $FCSD_Y$ | PC 1 | 0.800 | 0.800 | 0.690 |
|  | PC 2 | 0.157 | 0.157 | 0.154 |
|  | PC 3 | 0.023 | 0.023 | 0.090 |
| $\text{FCSD}_\beta$ | PC 1 | 0.906 | 0.859 | 0.520 |
|  | PC 2 | 0.079 | 0.108 | 0.329 |
|  | PC 3 | 0.013 | 0.019 | 0.092 |
| $\overline{\text{FCSD}}_\alpha$ | PC 1 | 0.259 | 0.798 | 0.733 |
|  | PC 2 | 0.513 | 0.158 | 0.108 |
|  | PC 3 | 0.050 | 0.022 | 0.092 |

**Figure 4.**
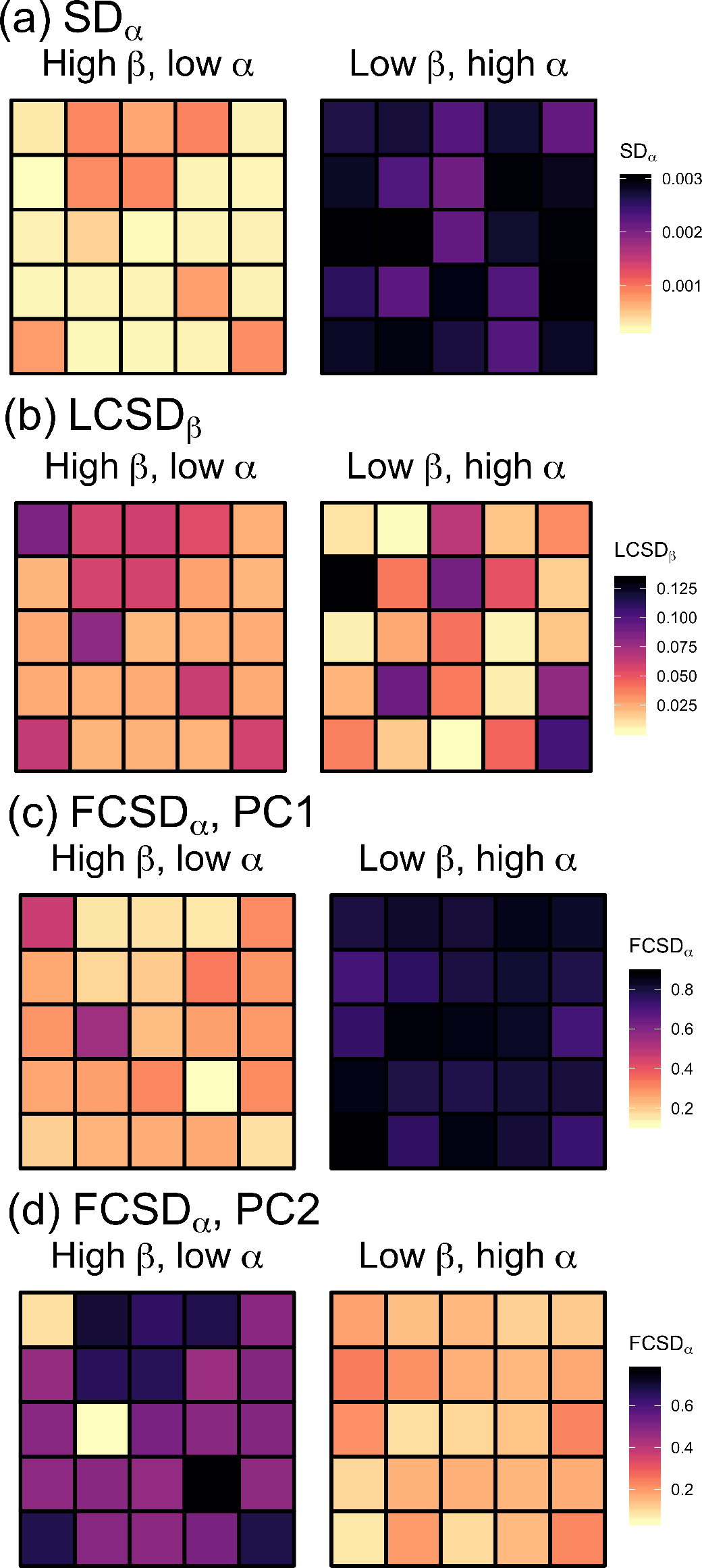
(a) Spectral *α*-diversity (SD_*α*_), (b) local contribution to spectral *β*-diversity (LCSD_*β*_), and (c-d) feature contribution to spectral *α*-diversity (FCSD_*α*_) for the first two spectral features (i.e. first two principal components of the brightness-normalized reflectance data) of each community in the two simulated regions. PC = principal component.

Next, we determined the local contributions of individual communities to spectral *β*-diversity (LCSD_*β*_). In the high *β*-diversity scenario (Fig. 4b, left panel), communities 12 and 21 (numbered as in Fig. 3) contributed the most to spectral *β*-diversity. These were the only two plots (out of 25) containing spectra of *Populus tremuloides*. In other words, these two plots had the most distinctive spectral composition compared to other communities. By contrast, in the low *β*-diversity scenario, community 16 was the most spectrally distinct community, something that could not be easily detected by examining this scenario visually (Fig. 3, right panel). As illustrated here, it is important to note that a region with low SD_*β*_ can still have individual communities showing high LCSD_*β*_ values, because LCSD_*β*_ values are *proportions* of SD_*β*_.

We then estimated the contributions of individual spectral features to spectral diversity (FCSD) for each scenario. For spectral *γ*-diversity (total variance of the region), FCSD_*γ*_ declined progressively from the first to the third PC (Table 1). As mentioned previously, FCSD_*γ*_ values are equal to the relative PCA eigenvalues of the spectral feature. Likewise, for spectral *β*-diversity, the contribution from the first to subsequent PCs decreased in both scenarios (Table 1). The relative contributions of individual spectral features to *β*-diversity were fairly similar in both regions, even though they differed considerably in spectral *β*-diversity. For spectral *α*-diversity, however, FCSD_*α*_ values differed noticeably among the two scenarios (Fig. 4c–d). In the low *α*-diversity scenario (Fig. 4c-d, left column), PC 2 contributed more strongly to the spectral *α*-diversity of most communities than PC 1, whereas the opposite was true for the high *α*-diversity scenario (Fig. 4c–d, right column). The FCSD_*α*_ values were not expected to decrease in a monotonic way since *α*-diversity is orthogonal to *γ*-diversity and the PCs are those of *γ*-, not of *α*-diversity.

We note that SS, SD and LCSD indices are exactly the same whether using the original spectral bands or all PCs, because PCA type-I scaling preserves the Euclidean distance among objects (e.g., image pixels in spectral space). Conversely, the equations developed in this paper hold and can be used directly with the original band data. However, FCSD values would change when using the original spectral bands instead of PCs, since this would then indicate the relative contributions of individual spectral bands (instead of PCs) to spectral diversity.

## CASE STUDY 2: NEON IMAGERY

Next, we applied our method for partitioning spectral diversity to imaging spectroscopy data collected by NEON’s Airborne Observation Platform (AOP; Kampe *et al.* 2010) over the Bartlett Experimental Forest (https://www.neonscience.org/field-sites/field-sites-map/BART). In this case study, we used a scene measuring 280 m (east-west) × 1000 m (north-south), acquired in August 2017. Spectral data were processed to surface reflectance and subsampled to 1-m pixel size by NEON. Our workflow is illustrated in Figure 2.

For spectral diversity calculations we selected a community (i.e. plot) size of 40 m × 40 m, which is the base plot size used by NEON. We used rarefaction to standardize the number of pixels per community used for analysis. We used a normalized difference vegetation index (NDVI) threshold of ≥0.8 to identify the minimum number of vegetated pixels across all plots in the image (termed *m*_min_), which was 1474 (= 92% of the 1600 pixels per community). We randomly selected *m*_min_ pixels per plot, and applied our spectral diversity partitioning approach to all selected pixels. The rarefaction was repeated 30 times and results were averaged across all 30 repeats (Fig. 2). Alternatively, one could take the median value instead of the mean if distributions are skewed.

Our analyses revealed that spectral *α*-diversity in this forested landscape accounted for 77% of the spectral *γ*-diversity, whereas *β*-diversity accounted for the remaining 23% (Table 1). In other words, there is considerably more spectral diversity within individual 40 m × 40 m communities than among communities in this forest. Figure 5 illustrates how spectral diversity is spatially structured. Two areas contribute strongly to spectral *β*-diversity (LCSD_*β*_, darker colours in Fig. 5). The tree communities in these areas are more spectrally dissimilar from the average community than communities with lower LCSD_*β*_ (lighter colours in Fig. 5).

**Figure 5.**
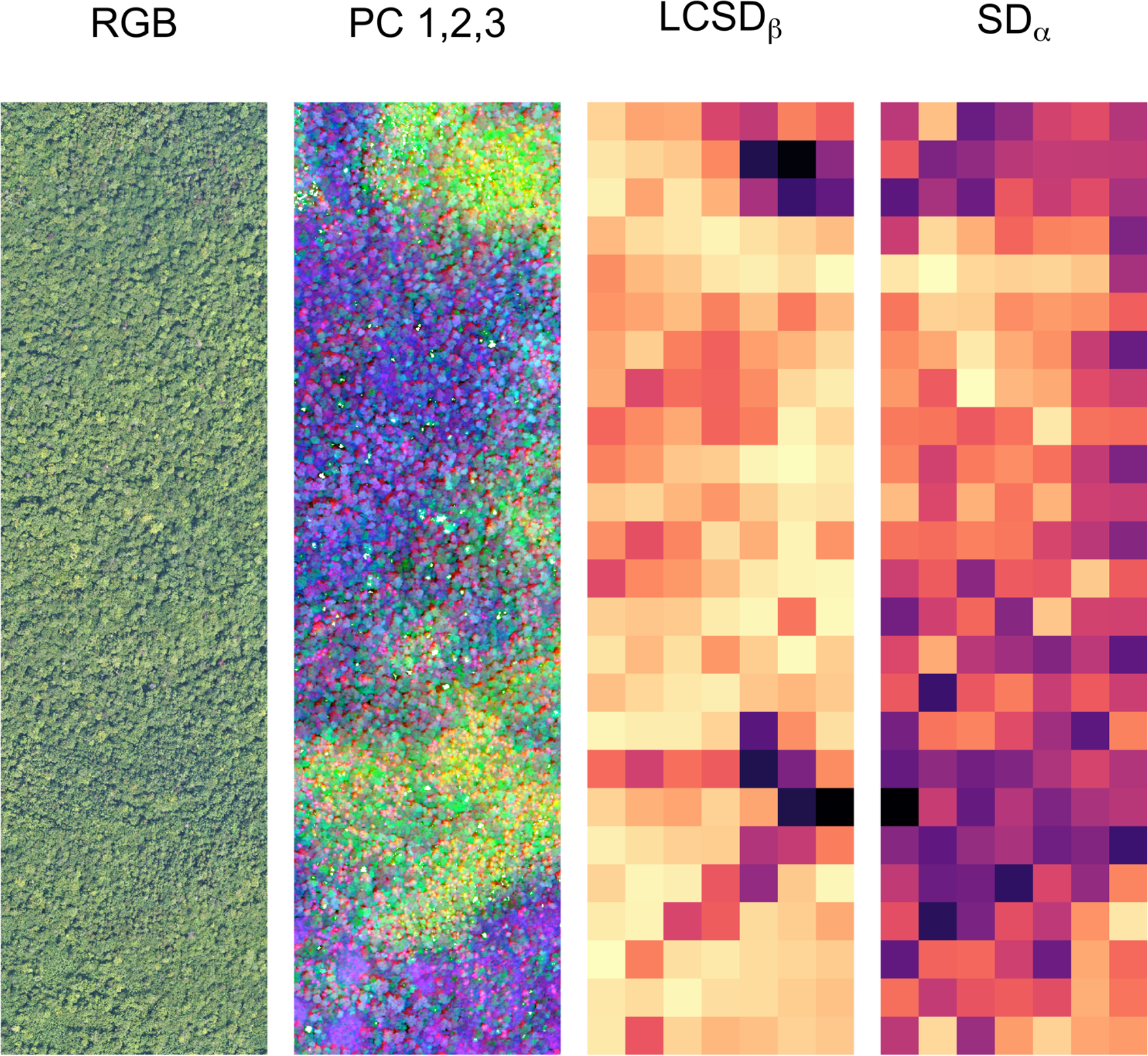
Partitioning spectral diversity using imaging spectroscopy data acquired by the National Ecological Observatory Network (NEON) over the Bartlett Experimental Forest site. From left to right: (1) true colour (red-green-blue, RGB) image with 0.1 m ground resolution, (2) false colour image with 1 m resolution based on the first three principal components (PCs) of the spectral image cube (PC1 = red, PC2 = green, PC3 = blue), (3) local contribution to spectral *β*-diversity (LCSD_*β*_), and (4) spectral α-diversity (SD_*α*_) of forest communities, each measuring 40 m × 40 m. For panels 3 and 4, light hues correspond to low, dark hues to high values of LCSD_*β*_ coefficients and SD_*α*_ values, respectively.

For completeness, we evaluated the effects of shadows and community size on spectral diversity calculations (Appendix S3). We found that removing shadows had little influence on spectral diversity (Figs. S4–S6) and that results remained remarkably stable for plots ranging from 20 m × 20 m (400 m^2^) to 140 m × 140 m (almost 2 ha) in size (Figs. S7–S9).

## DISCUSSION

In this paper, we proposed a new method for partitioning plant spectral *γ*-diversity (i.e. the spectral diversity of a region) into additive *α*= (within community) and *β*-diversity (among community) components. Our approach builds on a method for partitioning *β*-diversity initially designed for community data (Legendre & Cáceres 2013), adapts it to spectral data and, importantly, extends it to include *α, β and γ* components. Partitioning spectral diversity can bring new insights and generate new hypotheses about the origins and maintenance of plant spectral diversity across regions. For instance, high spectral *β*-diversity could result from turnover in plant species and/or functional trait composition across environmental gradients (e.g., soil properties, hydrology), whereas high spectral *α*-diversity might result from local biotic interactions among co-occurring plants (e.g., resource partitioning, conspecific negative density dependence). Mapping spectral indices such as LCSD_*β*_ and SD_*α*_ could be used as a biodiversity “discovery tool” to design targeted field sampling campaigns to test such hypotheses, e.g., by comparing community composition and diversity in areas with high and low LCSD_*β*_ and SD_*α*_ values, respectively (Fig. 5).

Partitioning spectral diversity allows the determination of the spectral features contributing most strongly to spectral *α*-, *β*- or *γ*-diversity (FCSD), which helps in understanding the underlying biological traits driving spectral diversity at different spatial scales. In our case studies, the spectral features were principal components (PCs), which are linear combinations of the original wavelength bands. As such, the individual contributions of all wavelength bands to each spectral feature can be retrieved. The bands, in turn, can be linked to specific plant properties, since the biophysical and biological causes of spectral variation across spectral regions and for specific absorption features of molecules are reasonably well understood (Gates *et al.* 1965; Curran 1989; Asner 1998; Kokaly *et al.* 2009; Ustin *et al.* 2009). Identifying the traits contributing most strongly to spectral *α*-diversity might inform us about how co-occurring species are partitioning resources at the local scale, whereas identifying the traits contributing most strongly to spectral *β*-diversity might reveal important mechanisms driving changes in community composition across environmental gradients.

Partitioning plant spectral diversity and mapping its spatial components has applications in biodiversity management. Indeed, managers often need to estimate the ecological value of different ecosystems over large regions, for example to prioritize conservation or restoration efforts. However, access to field data might be limited. Using imaging spectroscopy data, our approach of partitioning spectral diversity allows the identification of areas with high spectral *α*-diversity, which likely coincide with local “hotspots” of taxonomic and/or functional trait diversity. Further, high LCSD_*β*_ values indicates areas with rare spectral composition, i.e., containing communities that are most spectrally dissimilar from the average community within the region of interest. Given that species spectral dissimilarity is linked to their functional and phylogenetic dissimilarity (Schweiger *et al.* 2018), spectrally rare communities can be expected to have rare taxonomic and/or functional composition, either because they harbor uncommon species, or rare combinations of common species.

Our approach measures spectral variance directly (Fig. 2), which is in contrast to other studies that have prior to deriving biodiversity metrics first translated remotely-sensed spectra into plant species (e.g., Féret & Asner 2013), “spectral species” (Féret & Asner 2014), or plant functional traits (e.g., Dahlin *et al.* 2013; Schneider *et al.* 2017). While spectral diversity does not isolate any particular facet of plant biodiversity (e.g., taxonomic, chemical, structural), it integrates all of these facets (Schweiger *et al.* 2018; Appendix S1). From a practical perspective, casting spectral diversity as spectral variance depends on fewer user decisions compared to other approaches (e.g., selecting the number of clusters for classifying spectral species, selecting the plant traits and modelling approach to predict traits from spectra). This makes spectral diversity easily comparable across different regions. Therefore, maps of SD_*α*_ and LCSD_*β*_ could be ideal candidates for biodiversity products from remotely sensed spectral imagery.

### Comparison with other approaches

Much of the interest in measuring spectral diversity from remote sensing data stems from the spectral variation hypothesis (Palmer *et al.* 2002), stating that the spatial variation in spectral reflectance expresses overall variation of the environment. As areas of high environmental variation often harbour more species than areas with low environmental variation, spectral variation across space can potentially uncover botanically interesting areas (Palmer *et al.* 2002). However, spectral diversity has been predominantly used to investigate relationships between plant spectra and taxonomic units at the *α*- and *γ*-diversity scale, whereas the *β* component has received less attention (Rocchini *et al.* 2018).

Historically, Landsat satellites were instrumental for spurring large-scale biodiversity studies. Early sensors contained few spectral bands; thus, a large body of literature deals with using NDVI for predicting and mapping taxonomic diversity (Gould 2000; see review by Pettorelli *et al.* 2005). Recent advances in sensor technology, particularly increased spectral resolution, have led to a variety of approaches to calculate spectral *α*-diversity (Rocchini *et al.* 2010). This includes metrics such as the standard deviation or coefficient of variation of spectral indices (Oindo & Skidmore 2002), or spectral bands among pixels (Hall *et al.* 2010; Gholizadeh *et al.* 2018; Wang *et al.* 2018a), the convex hull volume of pixels in spectral feature space (Dahlin 2016), the mean distance of pixels from the spectral centroid (Rocchini et al. 2010), the number of spectrally distinct clusters or spectral species in ordination space (Féret & Asner 2014), and diversity metrics based on dissimilarity matrices among species spectra or image pixels (Schweiger *et al.* 2018). Of these, our method is most similar to the mean distance to the spectral centroid (Rocchini *et al.* 2010). The difference is that we square the individual distances to the spectral centroid; doing so allows us to partition sums of squares into additive components (Equation 9).

Fewer studies have considered spectral *β*-diversity (Rocchini *et al.* 2018). One approach for studying *β*-diversity using spectra has been to combine ordination scores of species inventories with spectral data in multivariate models to predict the positions of pixels with unknown species composition in species-ordination-space (Schmidtlein *et al.* 2007). This method and some of its variants (Rocchini *et al.* 2018) do not measure spectral *β*-diversity *per se*, but instead use spectra to estimate changes in community composition across the landscape. Rao’s quadratic entropy has been suggested as a measure of spectral *β*-diversity, based on the dissimilarity among image pixels within a moving window (Rocchini *et al.* 2018). However, a moving window approach expresses spectral *β*-diversity for many small sub-regions independently from one another and does not estimate the spectral *β*-diversity of the region as a whole. Another approach for studying spectral *β*-diversity has been to measure the pairwise dissimilarity in the composition of spectral species among mapping units, and to re-project those pairwise dissimilarities onto an RGB colour space (Féret & Asner 2014). This method yields a useful map showing changes in spectral composition across the region, similar to our mapping of the first three PCs in Figure 5, but it does not calculate spectral diversity.

### Methodological considerations

A number of methodological aspects should be considered before applying our approach to imaging spectroscopy data. These include: (1) the choice of a brightness normalization procedure, (2) whether all or a subset of the wavelength bands, or spectral features, should be used, (3) masking non-vegetated pixels, or not, (4) determining community size, and (5) deciding on the scaling type (i.e. type I or II; Legendre & Legendre 2012) if using PCA as a spectral feature extraction method. We discuss these methodological points in detail in Appendix S4.

## CONCLUSION

Plant spectral diversity is emerging as an integrator of chemical, structural, and taxonomic aspects of plant biodiversity, which can be remotely sensed (Cavender-Bares *et al.* 2017). Partitioning plant spectral diversity using our approach can help us to better understand and generate new hypotheses about the origins of, and the processes that drive, biodiversity variation across regions. Given the rapid and broad extent of current environmental changes, remote sensing of plant biodiversity over large regions is more important than ever (Turner 2014; Wang & Gamon 2019). Our approach can identify local *α*-diversity hotspots as well as unique areas contributing strongly to *β*-diversity – two central facets of biodiversity.

Our approach is timely since current technological developments in high-resolution UAV imaging spectroscopy will make this technology more accessible to ecologists in the coming years (Aasen *et al.* 2018; Arroyo-Mora *et al.* 2019). For example, the Canadian Airborne Biodiversity Observatory (www.caboscience.org) is developing UAV spectroscopy to understand how plant biodiversity is responding to major environmental changes across Canada. We anticipate that a growing number of ecologists will embrace this transformative technology for mapping plant biodiversity. In fact, a wealth of moderate-resolution imaging spectroscopy data are already freely available for a wide range of ecosystems across the United States as part of the NEON program (www.neonscience.org). Partitioning spectral diversity could become a useful tool for remotely sensing plant biodiversity from these new data sources.

## ACKNOWLEDGEMENTS

EL was supported by a Discovery Grant from the Natural Sciences and Engineering Research Council of Canada (NSERC; Grant numbers RGPIN-2014-06106, RGPIN-2019-04537). PL was supported by NSERC Discovery Grant number 7738. Teaching relief to EL that facilitated the writing of this manuscript was provided by the Canada Research Chair (CRC) in Plant Functional Biodiversity, and by Université de Montréal (UdeM) in support of a Discovery Frontiers grant from NSERC (Grant number 509190-2017) to EL to establish the Canadian Airborne Biodiversity Observatory (CABO). AKS was supported by a postdoctoral fellowship from CABO. EL wishes to thank colleagues from CABO and graduate students from the Laboratory of Plant Functional Ecology (LEFO) at UdeM for stimulating discussions about remote sensing of plant biodiversity, which helped to shape some of the ideas presented here.

