## Supporting Information for "Partitioning plant spectral diversity into alpha and beta components"

**Appendix S1** Discussion about the biological and ecological meaning of plant spectral diversity.

Plants express a suite of traits that have evolved over time and allow them to persist in a particular environment. In these environments the particular characteristics of individual plants within communities shape species interactions across trophic levels, with consequences for the functioning of the entire ecosystem, including the provisioning of nutrients, water and habitat, above- and belowground (Violle *et al.* 2014). The functional diversity of plant communities, which supports the diversity of life, can be calculated from plant functional dissimilarity within and across communities in different ways (e.g., Laliberté & Legendre 2010; Mouillot *et al.* 2013; Scheiner *et al.* 2017). However, the degree to which ecosystem consequences of biodiversity are captured by measures of functional diversity depends largely on the traits included. Although at the global scale the major axes of plant differentiation related to growth, survival and reproduction can be described using only six functional traits (adult plant height, specific stem density, leaf area, leaf mass per area, nitrogen content and diaspore mass; Díaz *et al.* 2016), this does not mean that other traits are irrelevant at smaller scale. But how many traits are needed to describe the variety of strategies and functions of plants in any particular environment?

Taxonomic diversity metrics are based on species identity, and by avoiding the selection of specific plant characteristics, they capture information about ecological variation among clades that cannot be resolved by functional traits alone (Clark 2010). However, plant clades (such as species) and individuals within these clades are not equally dissimilar from each other. Phylogenetic diversity metrics consider the general expectation, that the degree of dissimilarity among plants is proportional to the time since divergence from a common ancestor (Webb 2000). Close relatives are expected to be on average more functionally similar than distant relatives. However, the level of phylogenetic conservatism varies among traits and clades, and depends on the geographic context (Cadotte *et al.* 2017). For example, within a certain region, close relatives can be more dissimilar in their traits than expected due to character displacement, and distant relatives can more similar than expected due to convergence.

In contrast, spectral diversity neither depends on selecting functional traits *a priori* (although absorption features indicative of specific functional traits can be used), nor on the degree of phylogenetic conservatism of those traits (Schweiger *et al.* 2018) although there is evidence for plant spectral dissimilarity being tied to both functional and phylogenetic dissimilarity (Asner & Martin 2009; Cavender-Bares *et al.* 2016; McManus *et al.* 2016; Schweiger *et al.* 2018). Plants display themselves towards the sky in a myriad of ways, giving rise to specific spectral profiles that capture the environmental context and evolutionary legacy of individuals (Cavender-Bares *et al.* 2017). Chemical, structural, morphological and anatomical characteristics of leaves, as well as plant growth form and canopy architecture influence the reflectance, absorptance, transmittance of electromagnetic radiation (Gates *et al.* 1965; Curran 1989; Ustin & Gamon 2010). The degree of dissimilarity in plant life within a certain region that is captured by spectral diversity can expected to be similar to what is captured by functional diversity, when the traits included in the functional diversity measure are the most important drivers of spectral variation, phylogenetic diversity, when these traits are phylogenetically conserved, and taxonomic diversity, when the plant clades within the region are spectrally distinct. In addition, spectral diversity also captures important information about the vertical structure of the canopy, especially in the near-infrared range where scattering of photons within canopies dominate the spectral response (Asner 1998). Spectral diversity, the variation of plant spectra across space, thus integratively captures important aspects of the variability of plant phenotypes within a certain geographic area (Ustin & Gamon 2010; Schweiger *et al.* 2018).

**Appendix S2** Details about leaf spectra used in the simulation study.

Leaf spectra were measured with a portable field spectrometer (ASD FieldSpec 4, Malvern Panalytical, Cambridge, UK), covering the wavelength range from 350 nm to 2500 nm and an integrating sphere with internal light source (ASD RTS-3ZC, Malvern Panalytical, Cambridge, UK) in the summer of 2017, following the leaf spectroscopy protocol from the Carnegie Spectranomics project (https://cao.carnegiescience.edu/spectranomics-protocols), with some modifications (Laliberté & Soffer 2018). The tree species sampled (with five individual plants per species, each representing the average spectrum from six mature leaves) were *Betula alleghaniensis* Britton (yellow birch), *Populus deltoides* W. Bartram ex Marshall subsp*. deltoides* Marsh (eastern cottonwood), and *Populus tremuloides* Michaux (trembling aspen). Processing of spectra consisted of applying a third-order Savitzky-Golay filter (length = 55) to reduce noise, reducing spectral resolution from 1 nm to 10 nm wide to reduce the number of bands, trimming the spectra between 410 and 2400 nm to remove regions with low signal-to-noise, and brightness-normalizing spectra. This vector normalization emphasizes differences in the shape of spectra as opposed to differences in amplitude (i.e. albedo or brightness). The R code to perform the analyses is available online (https://github.com/elaliberte/specdiv).

**Appendix S3** Effects of shadows and community size on results of the NEON case study.

We evaluated the effects of shadows in the image on our results. To do so, we used the LiDAR-derived digital surface model (DSM) for our study area (Bartlett Experimental Forest) available from NEON and generated a shade mask, based on the solar zenith and azimuth angle at the time of image acquisition (Fig. S4). Overall, the presence of shadows had only minor effects on spectral diversity metrics in this example. Results were very similar (Figs. S5–S6, Pearson correlation coefficients $r > 0.93$), although $SD_\alpha$ values were slightly lower when shaded pixels were removed (Fig. S6).

We also determined how community or plot size impacted the results, using plot sizes ranging from 2 m × 2 m (i.e. 4 pixels) to 140 m x 140 m (i.e. 19 600 pixels). As expected, in small plots almost all spectral $\gamma$-diversity was expressed as $\beta$-diversity (Fig. S7), because when plots are approximately the size of an average canopy tree crown, spectral $\alpha$-diversity primarily reflects intra-individual spectral variability, which is comparatively small. The proportion of spectral $\gamma$-diversity expressed as $\beta$-diversity declined rapidly until plot sizes around 20 m × 20 m, after which the relative importance of $\beta$ vs $\alpha$ components stabilized (Fig. S7). In other words, increasing plot size had only minor effects on the partitioning of spectral diversity once plots reached a size large enough to contain several individual canopy trees (e.g. 20 m × 20 m, Figs. S7–S9).

**Appendix S4** Methodological considerations.

A number of methodological aspects are worth considering before applying our approach to imaging spectroscopy data. First, we used brightness normalization to emphasize differences in the shape of spectra rather than in albedo (Feilhauer *et al.* 2010); other methods (e.g., continuum removal) could also be used. Also, it could be argued that albedo differences might represent diversity in canopy structure to some degree (e.g., higher canopy roughness leading to more shaded areas), but our view is that it is best to remove differences in brightness, since these differences can be due to varying illumination and sensor viewing geometry, which are non-biological sources of spectral variation (Feilhauer *et al.* 2010; Féret & Asner 2014).

In our two case studies, we used PCA for extracting a set of uncorrelated spectral features, but we note that many other spectral feature extraction methods have been proposed in the literature (e.g., Bruce & and 2002). Alternatively, one may decide to use the original wavelength bands for spectral diversity calculation, but as we showed in our second case study, extracting spectral features (PCs in our case) can be useful for reducing data dimensionality and for removing artefacts that can arise during image acquisition and processing. In our case, it was straightforward to visually separate PCs containing biologically meaningful information (PCs 1-5) from noise (PCs 6-17, Fig. S3). However, developing automated methods for selecting spectral features would be one area warranting further research (Féret & Asner 2014).

Removing non-foliage pixels (e.g., soil), as we have done in our NEON case study by applying a NDVI threshold, can significantly improve the relationship between spectral diversity and taxonomic diversity of the top canopy layer (Gholizadeh *et al.* 2018). It also makes it compatible with plot-level vegetation data where canopy cover is used as the raw data from which taxonomic diversity is calculated. On the other hand, non-vegetated pixels containing soil, shadow, or rock might harbour ecosystem elements that are important for the diversity and abundance of understory plants or other trophic levels. As such, ecological knowledge should guide whether or not those pixels should be included in measures of spectral diversity. In our NEON case study, we found that masking crown shadows affected our results only marginally. However, removing poorly illuminated pixels can be important when imagery is acquired at lower sun angles (e.g., in the morning, in the afternoon, or at high latitudes), or when tall objects cast shadows on the vegetation layer under study (e.g. trees or buildings in grasslands). Differences in topography can be another source of variation in illumination requiring special consideration, such as splitting a region of interest into sub-regions based on slope and/or aspect.

One important user decision concerns the size of the community. At one extreme, if each community is only composed of one pixel, $\gamma$ diversity will be equal to $\beta$, and $\alpha$ will be zero. At the other extreme, if the community comprises the entire region, then $\gamma$ diversity will be equal to $\alpha$ and $\beta$ will be indeterminate. Obviously, reasonable community size parameters for real applications lie somewhere in between. Our analysis of NEON imagery showed that the proportion of spectral $\gamma$-diversity represented as $\beta$ vs. $\alpha$ remained remarkably stable for plots ranging from 20 m × 20 m (400 $m^2$) to 140 m x 140 m (almost 2 ha) in size. This indicates that our method for partitioning spectral diversity picks up variation in spectral community composition consistently, given that plots are large enough to contain a certain number of individuals. In our case, visual examination of aerial photographs shows that 400 $m^2$ plots contained around 15 individual canopy trees. Further studies are needed to investigate if this represents a generalizable threshold for the number of individuals per community, after which the fraction of $\beta$:$\alpha$ diversity stabilizes. We note that 400 $m^2$ is a plot size that is often used for tree surveys in temperate forests (e.g., Canada's National Forest Inventory 2008). Aligning community size for spectral diversity calculations to the plot size used in field surveys seems justifiable if it facilitates the interpretation of results.

Analyses of our two case studies were done using spectral features (i.e. PCs) obtained using PCA type I-scaling. Type-I scaling in PCA preserves the Euclidean distances among pixels in spectral feature space, and the PCs, which are used to compute spectral diversity, are uncorrelated with one another by definition. However, the original spectral data from which PCs are obtained typically show a high degree of correlation, i.e., neighbouring wavelength bands often expressing similar information. It follows that the first few spectral features (i.e. PCs) to which a large number of wavelength bands contribute will represent a higher proportion of the total variance of the spectral matrix than the use of the same number of selected original wavelength bands. The first PCs will therefore be weighted more strongly in spectral diversity analyses. For example, the near infrared (approximately 700–1000 nm) region contains many bands that together express differences in leaf internal structure and/or canopy architecture (Gates *et al.* 1965; Asner 1998). Reflectance in this region is generally high and also highly variable (e.g., Asner *et al.* 2014) because leaves absorb little radiation in this region (Merzlyak *et al.* 2002). By contrast, phenolics (e.g., tannins) play important biological roles in terms of foliar defense against herbivores (Chauvin *et al.* 2018), but show only a narrow absorption feature centered around 1660 nm in fresh leaves (Kokaly & Skidmore 2015) and therefore contribute little to the total spectral variance. As such, one could argue that certain dimensions of plant spectral variation are ecologically important even if they only represent a small proportion of the overall variance. If so, a case can be made for using PCA with type-II scaling, which preserves the Mahalanobis distance among objects (Legendre & Legendre 2012) and accounts for the covariance among wavelengths bands. In practice, PCA with type-II scaling weighs each spectral feature (i.e. each PC) equally when calculating spectral $\gamma$-diversity because the PCs are rescaled to equal variances of 1. Consequently, it becomes critical to select only biologically meaningful spectral features to avoid over-emphasizing sources of variation that are not biologically meaningful, including atmospheric influences or instrument noise (e.g., PCs 6 to 17 in our NEON example). Figure S10 shows the results of spectral variation partitioning when type-II scaling is used for the NEON data. In contrast to type-I scaling, the spectral features contributing most strongly to $\beta$-diversity differ from those contributing most strongly to $\alpha$-diversity. The maps of $LCSD_\beta$ and $SD_\alpha$ based on type-II scaling (Fig. S10) also differ qualitatively from those based on type-I scaling (Fig. 5), but differences are relatively small overall. We believe that choosing the type of scaling in PCA deserves further attention; until then, we recommend using PCA with type-I scaling by default, and selecting PCs or individual spectral bands of biological importance when type-II scaling is used.

Although measuring spectral variance does not require any field data, we want to emphasize that mapping spectral diversity and its components should be viewed as a first step in regional-level biodiversity assessments. Mapping spectral diversity components (e.g., Fig. 5) highlights areas that are spectrally diverse and/or unique, and therefore warrant our attention. However, the ecological interpretation of spectral diversity and distinctiveness is still in its infancy and depends on field data, although automated interpretation might be possible in the future, e.g., through machine learning (Brodrick *et al.* 2019). One way around the less well-understood aspects of spectral diversity would be focusing only on spectral regions that are tied to chemical components with well-known absorption features, e.g., pigments, phenolics, and using only the wavelength bands at these absorption features for calculating spectral diversity. However, the absorption features of most plant traits are subtle and often overlap one another, and leaf and canopy structure affect large portions of the spectrum (Asner 1998; Kokaly *et al.* 2009; Ollinger 2010). Although certainly useful for investigating specific questions, we believe that by limiting analyses to a few well-understood spectral features, one misses out on capturing plant characteristics tied to the spectral response that are important facets of plant biodiversity (Cavender-Bares *et al.* 2017). In addition, we note that our approach for calculating the contribution of each individual wavelength band or spectral features (which are PCA linear combinations of bands) to spectral diversity (LCSD) allows one to establish a direct link to functional trait models, e.g. via partial least squares regression (PLSR) model coefficients, thereby enabling the identification of the contributions of specific plant traits to spectral diversity.

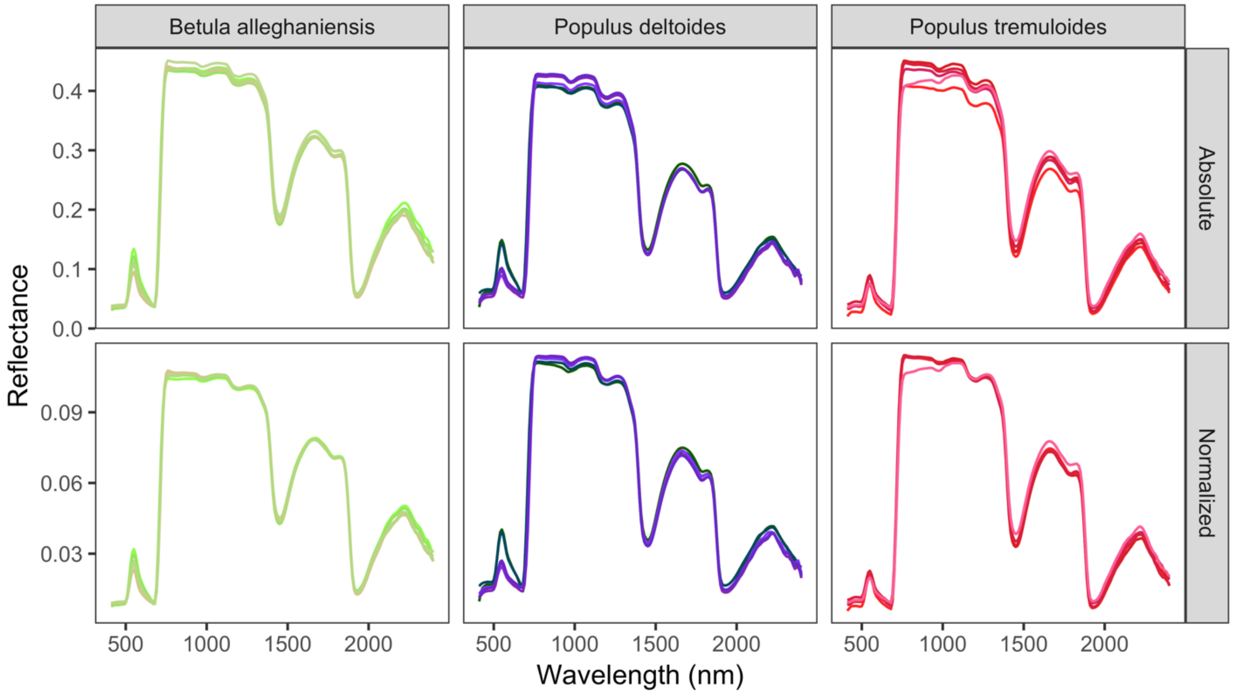


**Figure S1** Foliar reflectance spectra for 15 individual trees from three species. The top row shows absolute reflectance data, the bottom row shows brightness-normalized spectra. Comparing the bottom and top row shows how normalized spectra emphasize differences in shape while suppressing difference in brightness. Processing of spectra consisted of applying a third-order Savitzky-Golay filter (length = 55) to reduce noise, reducing spectral resolution from 1 nm to 10 nm to reduce the number of bands, and trimming the spectra between 410 and 2400 nm to remove regions with low signal-to-noise ratio. The colour of each spectrum is based on the PCA ordination shown in Figure S2.

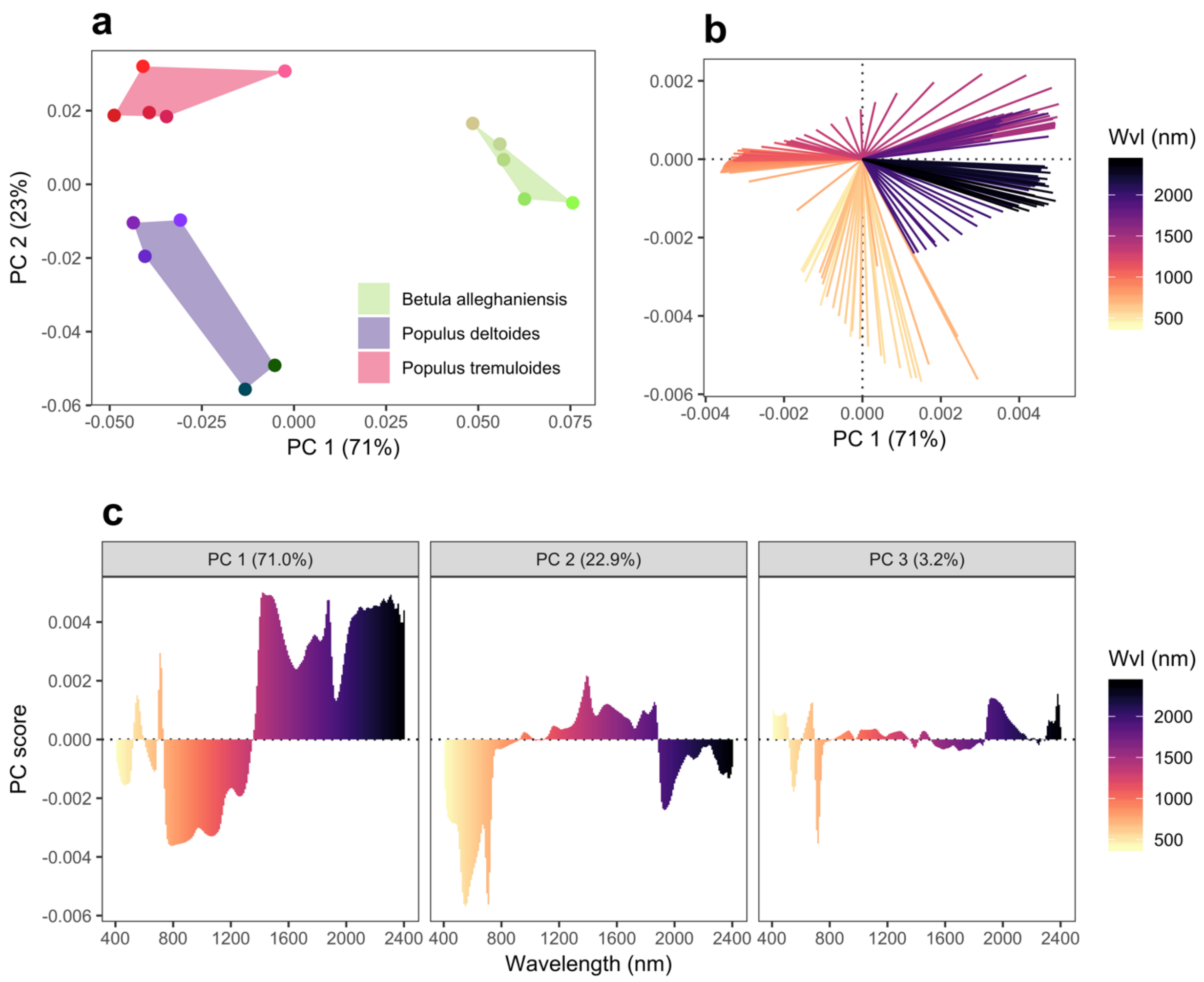


**Figure S2** Principal component analysis (PCA) of the brightness-normalized spectra in Figure S1. (**a**) Distance biplot (i.e. type-I scaling) showing the 15 spectra from three species (five spectra per species). Polygons are convex hulls linking the spectra for each species. Colours were set by mapping the scores of the first three principal components (PC) for each spectrum on a red-green-blue (RGB) scale (PC 1 = green, PC 2 = red, PC 3 = blue) to maximize contrast. Distances between spectra in the biplot approximate their Euclidean distances in feature space. (**b**) Correlation biplot (i.e. type-II scaling) showing all vectors (i.e. 10-nm wavelength bands). Angles between vectors approximate their correlations (0° = perfect positive correlation, 180° = perfect negative correlation, 90° = no correlation). The biplot shows the high degree of spectral autocorrelation in the data. (**c**) PC scores for each vector (type-II scaling as in **b**). Scores for each band represent their contribution to the formation of the first three PCs. Wvl = wavelength.

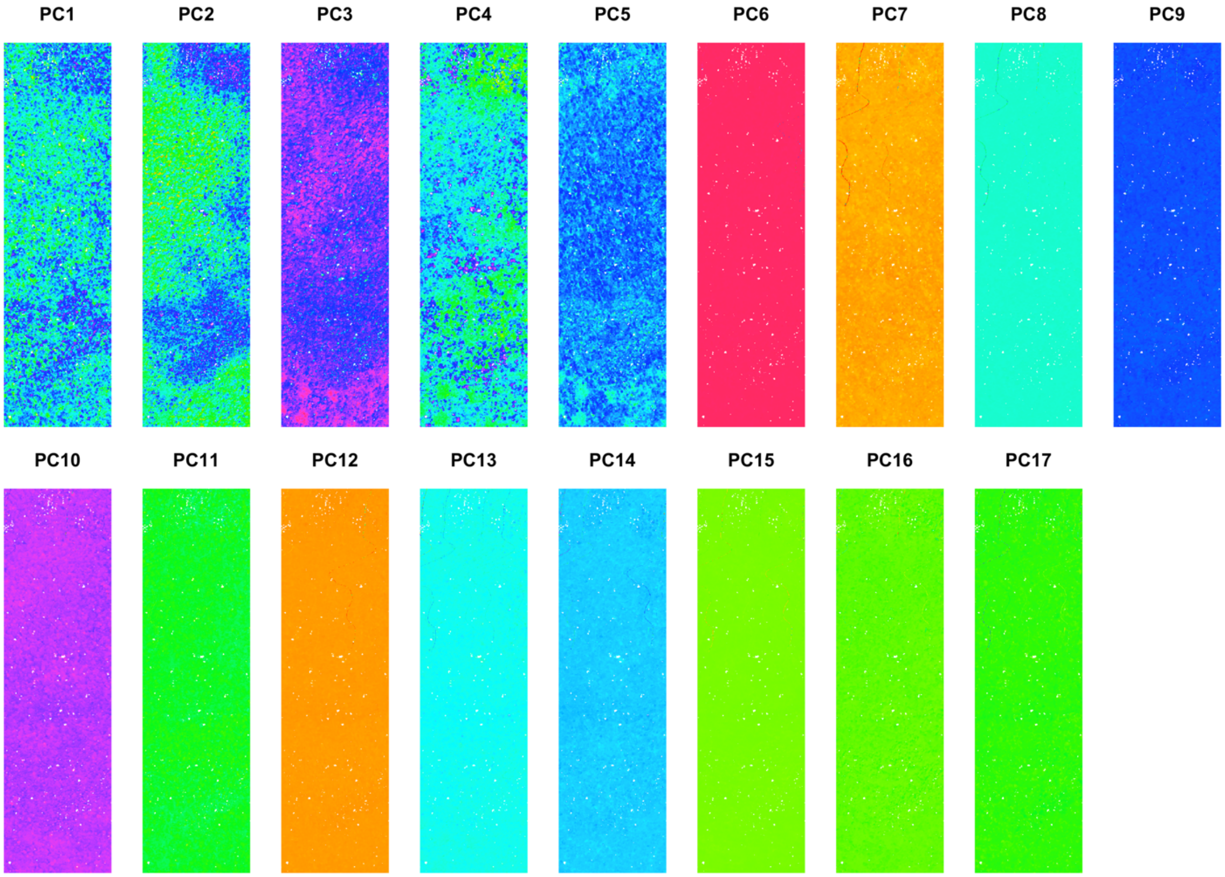


**Figure S3** The first 17 principal components (PCA type-I scaling) together representing >99% of the overall spectral variation among pixels for the NEON imagery data. Principal components 1–5 were retained for spectral diversity measurements since they showed biologically meaningful spatial patterns, whereas PCs 6–17 expressed image artefacts and were therefore excluded.

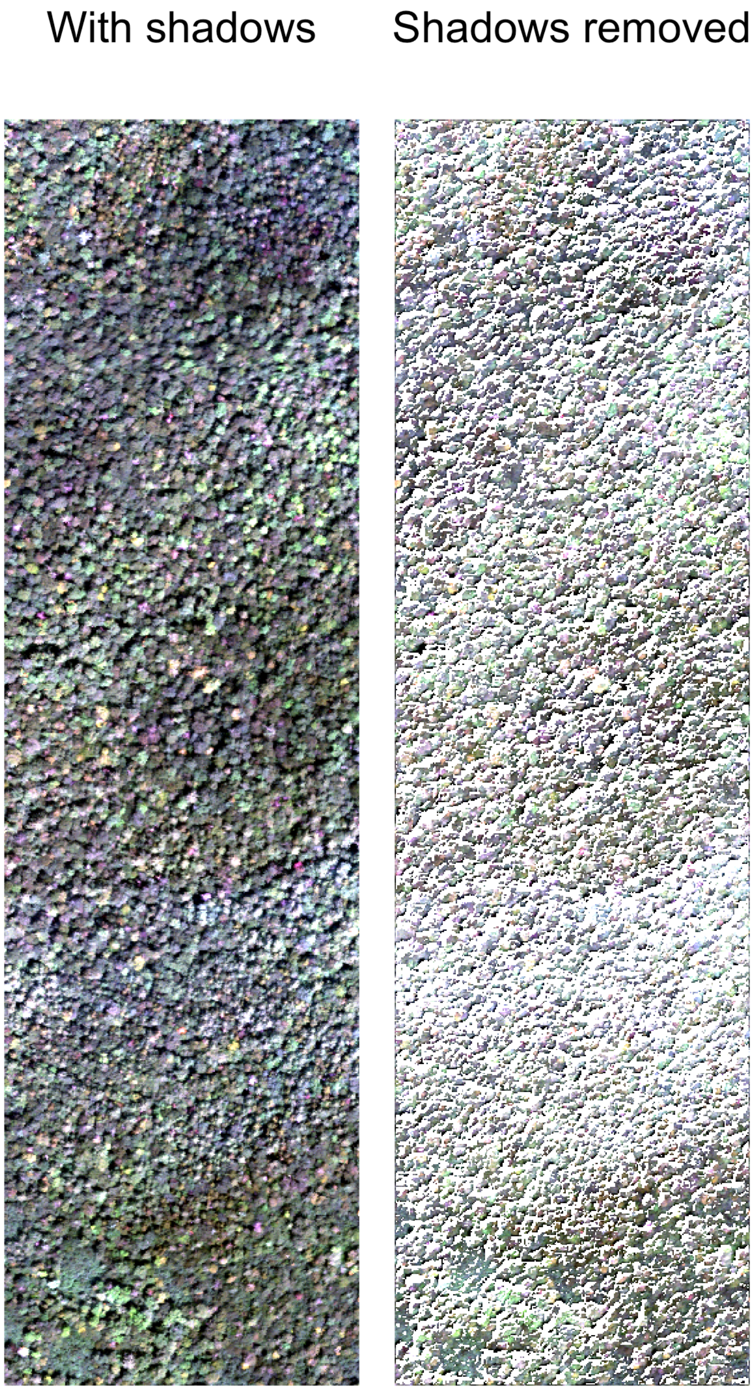


**Figure S4** True color RGB composite of the NEON imagery (left panel) showing canopy shadows and (right panel) after shadows were removed, using a shade mask derived from the digital surface model (DSM).

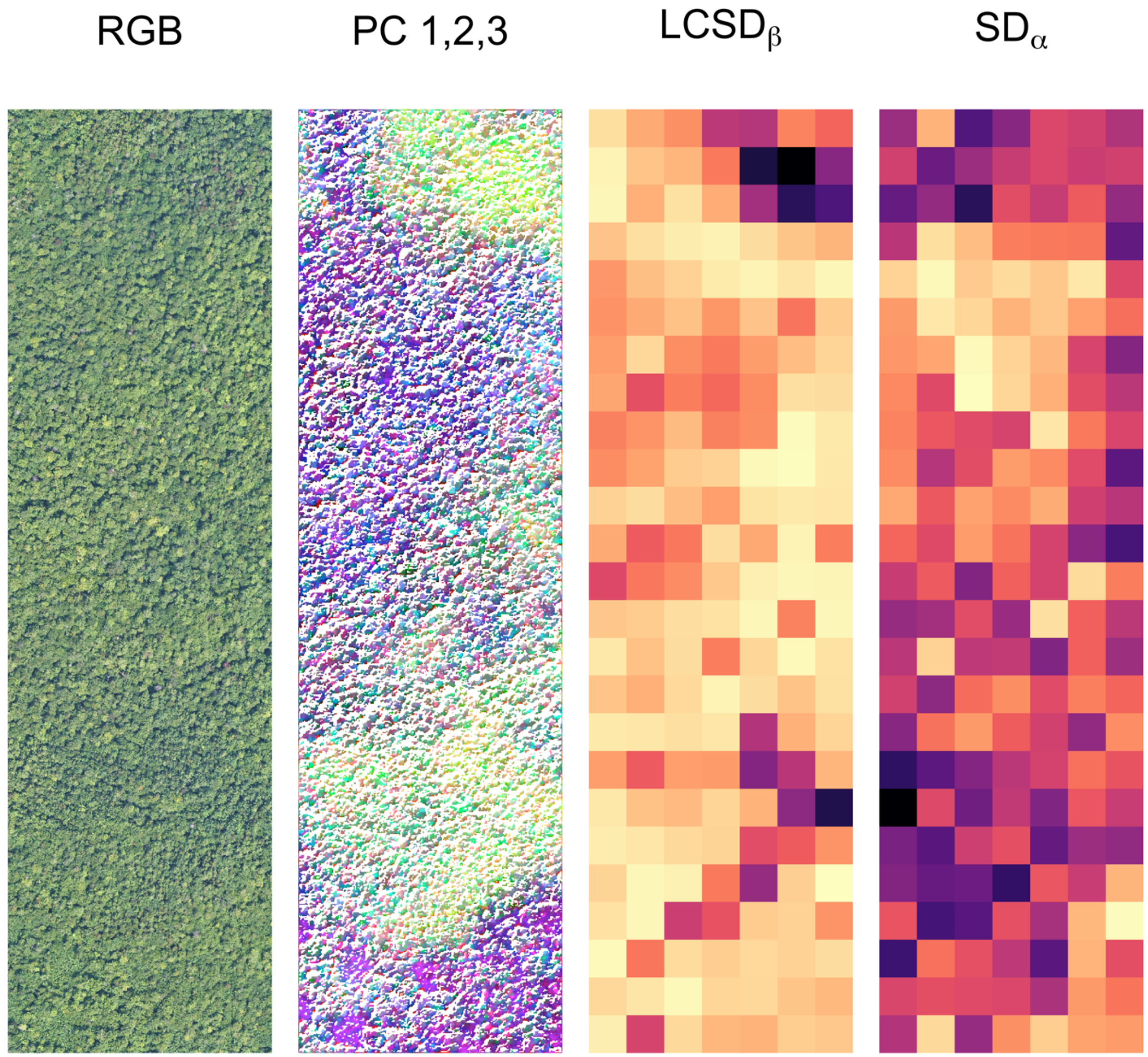


**Figure S5** Repeat of the analyses shown in Figure 5, but with shadows removed using the shade mask shown in Figure S4.

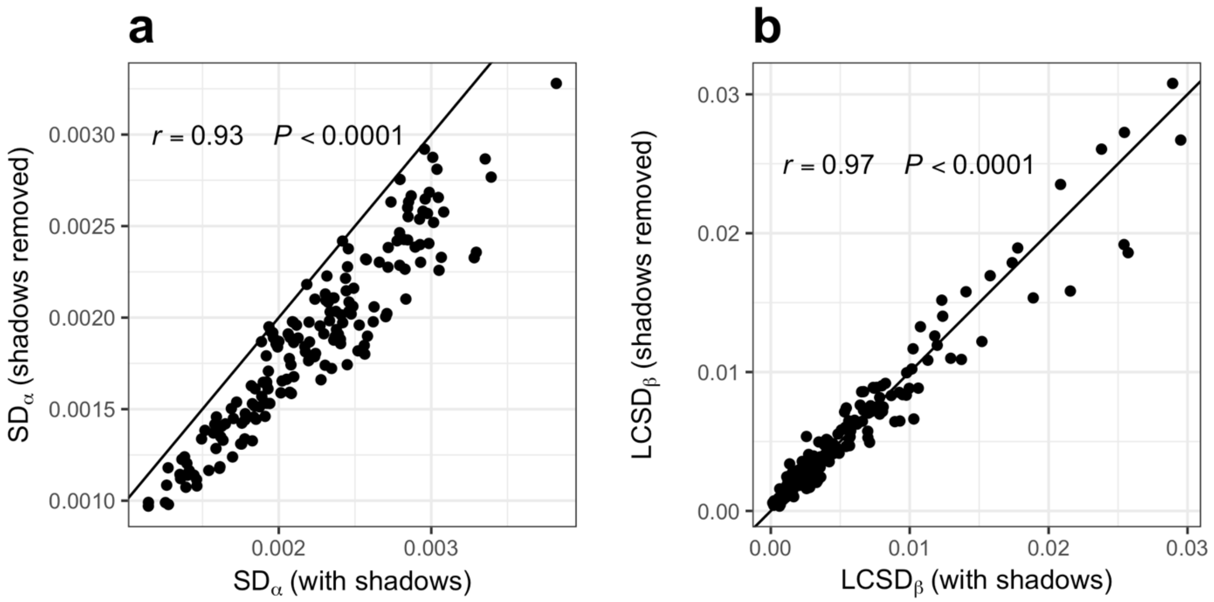


**Figure S6** Pairwise relationships between values for (**a**) $SD_\alpha$ and (**b**) $LCSD_\beta$ obtained from images with or without shadows. The 1:1 lines are shown (solid black lines).

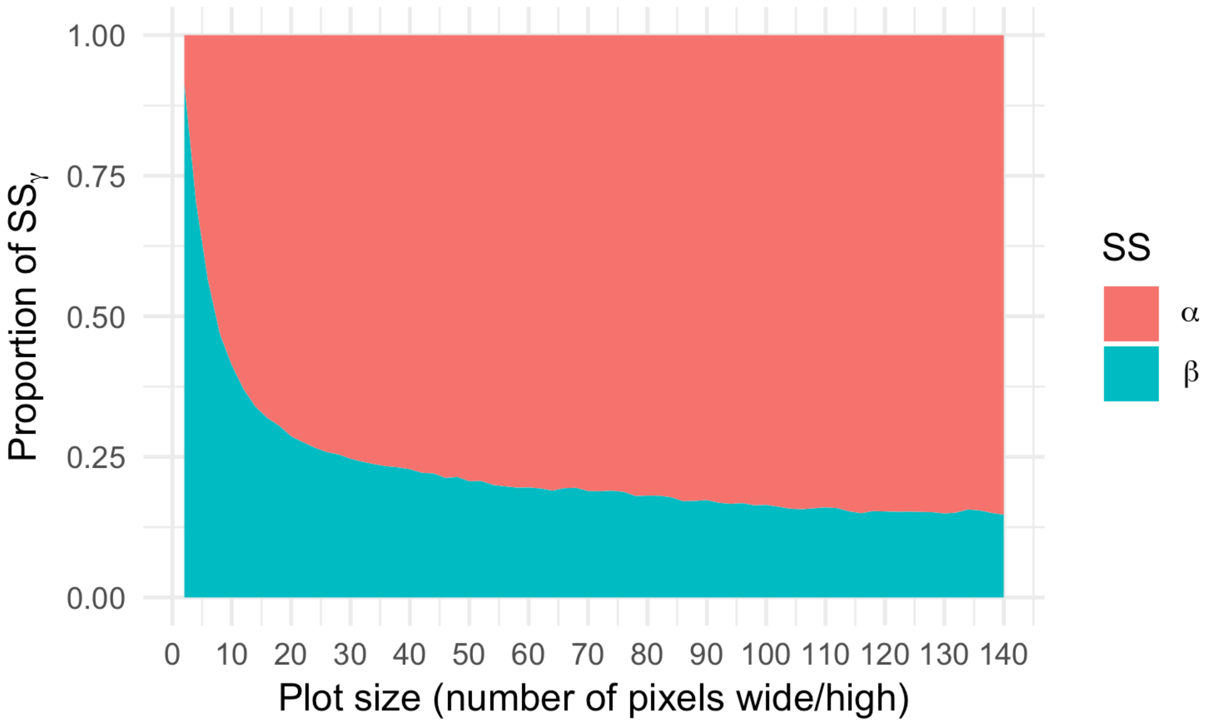


**Figure S7** Proportion of $SS_\gamma$ expressed as $SS_\beta$ vs. $SS_\alpha$ as a function of plot size.

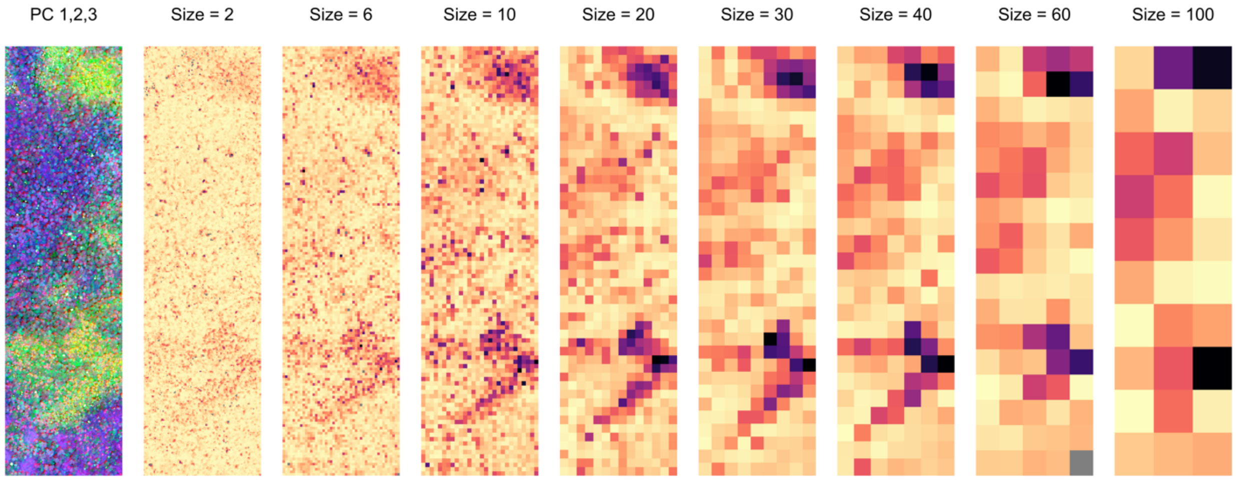


**Figure S8** Left panel: RGB false color composite showing the first three PCs of the brightness-normalized reflectance data. Other panels: $LCSD_{\beta}$ values as plot size increases from 2 pixels wide/high to 100 pixels wide/high.

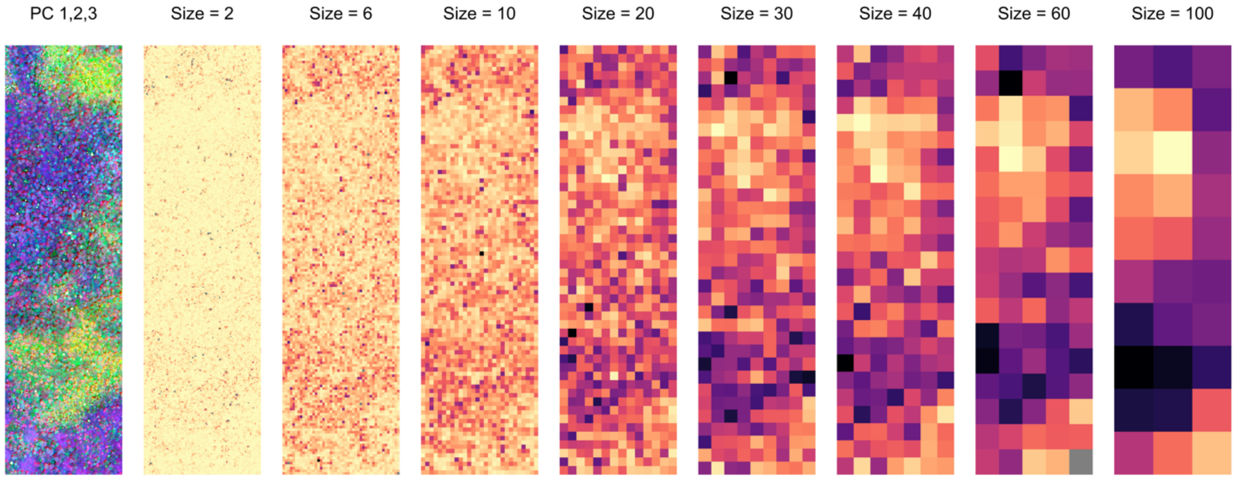


**Figure S9** Left panel: RGB false color composite showing the first three PCs of the brightness-normalized reflectance data. Other panels: $SD_{\alpha}$ values as plot size increases from 2 pixels wide/high to 100 pixels wide/high.

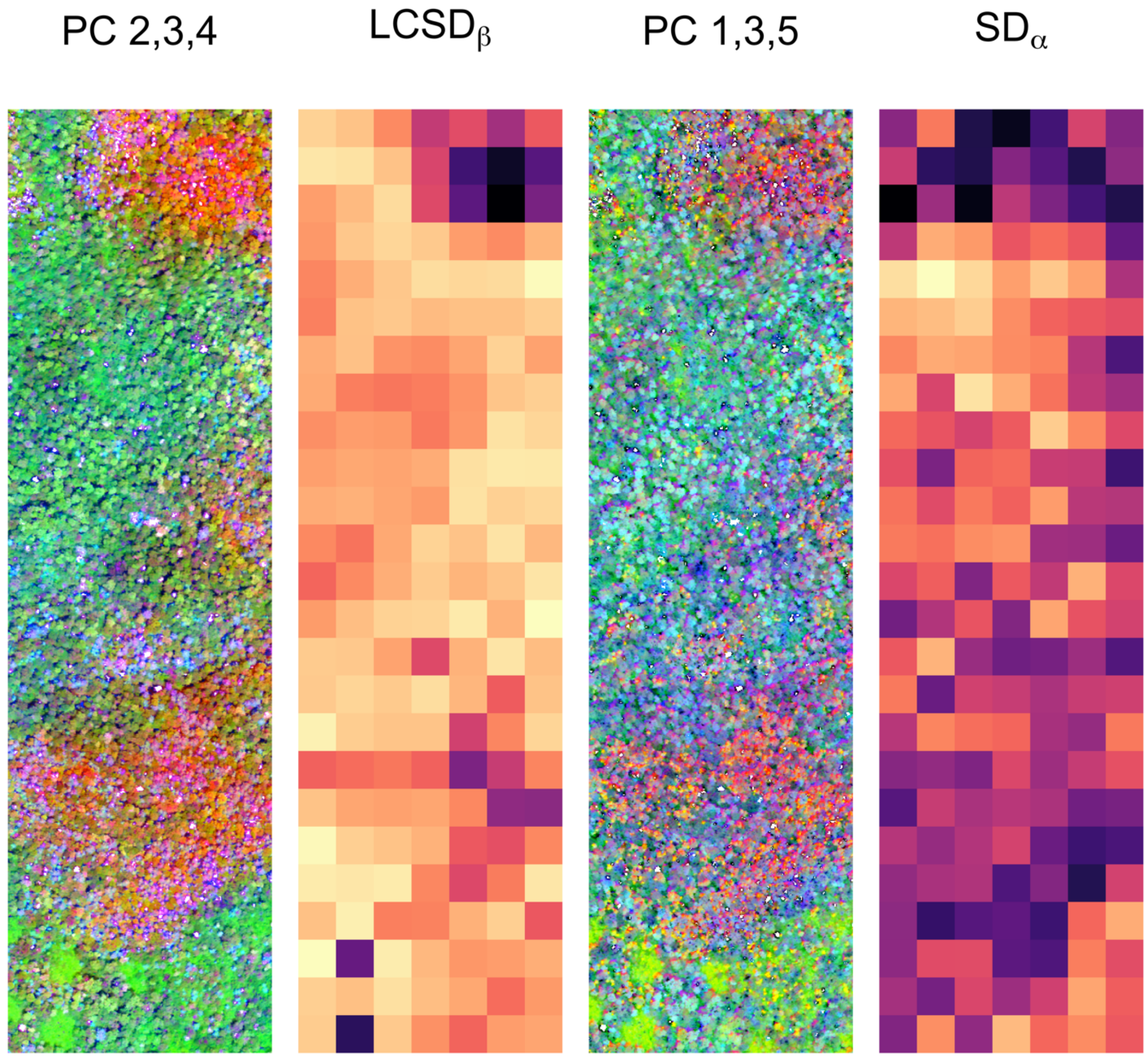


**Figure S10** Partitioning of spectral diversity at the NEON site Bartlett Forest using spectral features (i.e. principal components, or PCs) calculated using PCA type-II scaling, which preserves the Mahalanobis distance among pixels and therefore accounts for spectral autocorrelation in the original data (i.e. wavelength bands). The first panel on the left shows the three PCs that contribute most strongly to spectral $\beta$-diversity (PC2 = red, PC3, = green, PC4 =

blue), the third panels shows the three PCs which contribute most strongly to spectral $\bar{\alpha}$-diversity (PC1 = red, PC3 = green, PC5 = blue).
